# Landscape, complexity and regulation of a filamentous fungal transcriptome

**DOI:** 10.1101/2021.11.08.467853

**Authors:** Ping Lu, Daipeng Chen, Zhaomei Qi, Haoming Wang, Yitong Chen, Qinhu Wang, Cong Jiang, Huiquan Liu, Jin-Rong Xu

## Abstract

Alternative splicing (AS) and alternative polyadenylation (APA) of pre-mRNAs contribute greatly to transcriptome complexity and gene expression regulation in higher eukaryotes. Their biological impact in filamentous fungi, however, has been poorly studied. Here we combine PacBio Isoform Sequencing and strand-specific RNA-Seq of multiple tissues together with mutant characterization to reveal the landscape, complexity and regulation of AS and APA in the filamentous plant pathogenic fungus *Fusarium graminearum*. We updated the reference genome and generated a comprehensive annotation comprising 51,617 transcript isoforms from 17,189 genes. Majority of the transcripts represent novel isoforms, including 2,998 undiscovered protein-coding genes. In total, 42.7% of multi-exonic genes and 64.8% of genes have AS and APA isoforms, respectively, suggesting AS and APA increase previously unrecognized transcriptome complexity in fungi. Nonsense-mediated mRNA decay factor FgUPF1 may not degrade AS transcripts with premature-stop codons but regulate ribosome biogenesis. Distal polyadenylation sites have a strong signal but proximal polyadenylation isoforms are high expressed. The core 3’-end processing factors FgRNA15, FgHRP1, and FgFIP1 play important roles in promoting proximal polyadenylation site usage and also intron splicing. Genome-wide increase in the abundance of transcripts with retained introns and long 3’-UTRs and downregulation of the spliceosomal and 3’-end processing factors are found in older tissues and quiescent conidia, indicating that intron retention and 3’-UTR lengthening may be a transcriptional signature of aging and dormancy in fungi. Overall, our study generates a comprehensive full-length transcript annotation for *F. graminearum* and provides new insights into the complexity and regulation of transcriptome in filamentous fungi.

## INTRODUCTIONS

The Kingdom Fungi encompasses an enormous diversity of species with varied morphologies, ecologies, and life cycle strategies, including unicellular yeasts, multicellular filamentous fungi, and many plant and animal pathogens. Recent studies have revealed that the plant and animal transcriptomes are highly complex and that the post-transcriptional regulatory process of pre-mRNAs, including alternative splicing (AS), alternative polyadenylation (APA), and nonsense-mediated mRNA decay (NMD), contribute significantly to enhance functional diversity and regulate gene expression in multiple ways (Tian and Manley 2017; Chaudhary et al. 2019; Kishor et al. 2019). In fungi, however, the biological impact of these post-transcriptional regulatory mechanisms has not been well studied except in the budding yeast *Saccharomyces cerevisiae*, which has only approximately 300 intron-containing genes and a simple saprophytic life cycle without elaborate asexual or sexual structures/fruiting bodies. Although AS events have been analyzed by Illumina short-read RNA sequencing (RNA-Seq) analysis in *Trichoderma longibrachiatum, Verticillium dahliae*, and a few other species (Zhao et al. 2013; Xie et al. 2015; Gehrmann et al. 2016; Dong et al. 2017; Jin et al. 2017; Liu et al. 2020; Ibrahim et al. 2021), there is no comprehensive analysis on splice isoforms in filamentous fungi. Besides short-read RNA-Seq data being unsuitable for accurately reconstructing full-length splice isoforms, most of the published fungal RNA-Seq data are not strand-specific and overlapping transcripts between adjacent genes is widespread in fungi due to high gene density (Testa et al. 2015), which making transcript assembly problematic. Knowledge of full-length transcript isoforms is necessary to deduce encoded proteins and assess the roles of splice isoforms in gene regulation. Isoform sequencing (Iso-Seq) with PacBio single molecular real-time (SMRT) long-read technology offers a considerable advantage in characterizing transcriptome-wide full-length splice isoforms and post-transcriptional regulatory events without assembly. Although it has been widely used in plant and animal studies (www.pacb.com/applications/rna-sequencing/rna-sequencing-for-plant-and-animal-sciences/), to date Iso-Seq was only used for identifying polycistronic transcripts in a few species of mushroom forming fungi (Gordon et al. 2015) and reconstructing the full-length transcriptome of an entomopathogenic fungus *Ascosphaera apis* (Chen et al. 2020).

The filamentous ascomycete fungus *Fusarium graminearum* is the predominant causal agent of Fusarium head blight (FHB), one of the most devastating diseases on cereal crops (e.g., wheat, barley and corn) worldwide. The disease often leads to significant losses of grain yield and quality, and contamination with mycotoxins such as deoxynivalenol (DON) and zearalenone harmful to humans and animals (Goswami and Kistler 2004). In addition to FHB, *F. graminearum* causes other diseases in cereals during their life cycle including crown rot, root rot, and seedling blight (Zhou et al. 2018). Given its scientific/economic importance, *F. graminearum* has been listed as one of the top 10 most important fungal pathogens, ranking the fourth (Dean et al. 2012).

Although both conidia (asexual spores) and ascospores (sexual spores) of *F. graminearum* can initiate infection on the host, the ascospores discharged from perithecia (dark black fruiting bodies) are the primary inoculum of FHB (Shaner 2003). *F. graminearum* is homothallic and produces abundant perithecia relatively synchronously under laboratory conditions, making it an ideal system for studying sexual development. Recently, A-to-I mRNA editing, a novel fungal epigenetic phenomenon, was discovered to specifically occur during its sexual stage (Liu et al. 2016; Bian et al. 2019). Moreover, the relatively high rate of homologous recombination and ability to easily obtain homokaryotic transformants have made molecular studies in *F. graminearum* efficient and productive (Trail 2009; Kazan et al. 2012; Ma et al. 2013). To date, more than two thousand genes have been functionally characterized by targeted deletion or disruption in *F. graminearum*, including large-scale gene knockout projects with genes encoding transcription factors (Son et al. 2011), protein kinases (Wang et al. 2011), phosphatases (Yun et al. 2015), cytochrome P450 monooxygenases (Shin et al. 2017), putative peroxidases (Lee et al. 2018), ABC transporters (Yin et al. 2018), G protein-coupled receptors (Jiang et al. 2019), and orphan proteins (Jiang et al. 2020).

Comparative and functional genomic studies rely on accurate genome assembly and annotation. *F. graminearum* is one of the earliest fungal plant pathogens with its genome sequenced. In fact, it was the third filamentous fungus with its genome sequence published (Cuomo et al. 2007). Owing to its small genome size and low repetitive sequence content, the reference genome assembly of *F. graminearum* strain PH-1 is of extremely high quality (King et al. 2015; King et al. 2017). Nevertheless, errors in the reference genome and annotation were often observed (Liu et al. 2016). Moreover, despite the annotation has been improved by multiple revisions (King et al. 2017), currently available gene models of *F. graminearum* are mostly derived from computational prediction, which is incomplete or sometimes inaccurate. Most gene models contain only coding sequences (CDS), missing 5’- and/or 3’-untranslated regions (UTRs). No splice isoform information is available even in the latest annotation. Furthermore, although transcriptome data have been accumulated with GeneChip or RNA-Seq analyses to determine gene expression profiles during the life/infection cycle of *F. graminearum* (Dash et al. 2012; Kazan and Gardiner 2018; Brauer et al. 2020), few studies have investigated the landscape and regulation of its transcriptome, especially in terms of splice isoforms and APA.

Here, we improved the reference genome and annotation of *F. graminearum* by using PacBio SMRT long-read sequencing and presented the first comprehensive analysis of a full-length transcriptome in a filamentous fungus. We characterized the landscape of AS, APA, lncRNA, and polycistronic transcripts and revealed their regulation in different cell types based on Iso-Seq and strand-specific RNA-Seq data together with mutant characterization. Overall, our study generated an updated reference genome and comprehensive reference set of transcript isoforms for *F. graminearum*, and provided new insights into the complexity and regulation of transcriptome in filamentous fungi.

## RESULTS

### Revising the reference genome of PH-1 by PacBio and Illumina sequencing

To improve the reference genome of *F. graminearum*, we generated 13 Gb (>300x) high-quality long-reads of PH-1 (YL lab stock) with PacBio Sequel platform (Supplemental Table S1) and assembled them into 10 contigs *de novo* using *Canu* software (Koren et al. 2017) (Fig. 1A). By comparing the PacBio assembly with the most recent public assembly RR1 (Ensembl Fungi) of PH-1, a total of 315 different regions/sites were identified. To determine which assembly is correct at these regions/sites, we re-sequenced the PH-1 lab stocks from three different labs (YL, ZJU, and MSU) by Illumina and obtained >90x high-quality short reads for each lab stock (Supplemental Table S2). The Illumina short-reads of the three PH-1 lab stocks and PacBio long-reads were aligned to the two genome assemblies, respectively. Based on the alignments, 7 mis-assembly regions caused by repeat sequences, 200 base errors, and 42 InDel errors in the RR1 assembly (Fig. 1A-C; Supplemental Table S3), which impacts the annotation of 74 protein-coding genes, were evidenced by at least two PH-1 lab stocks. We corrected all the errors in the RR1 assembly and generated an updated version of PH-1 assembly (named YL1).

**Figure 1.**
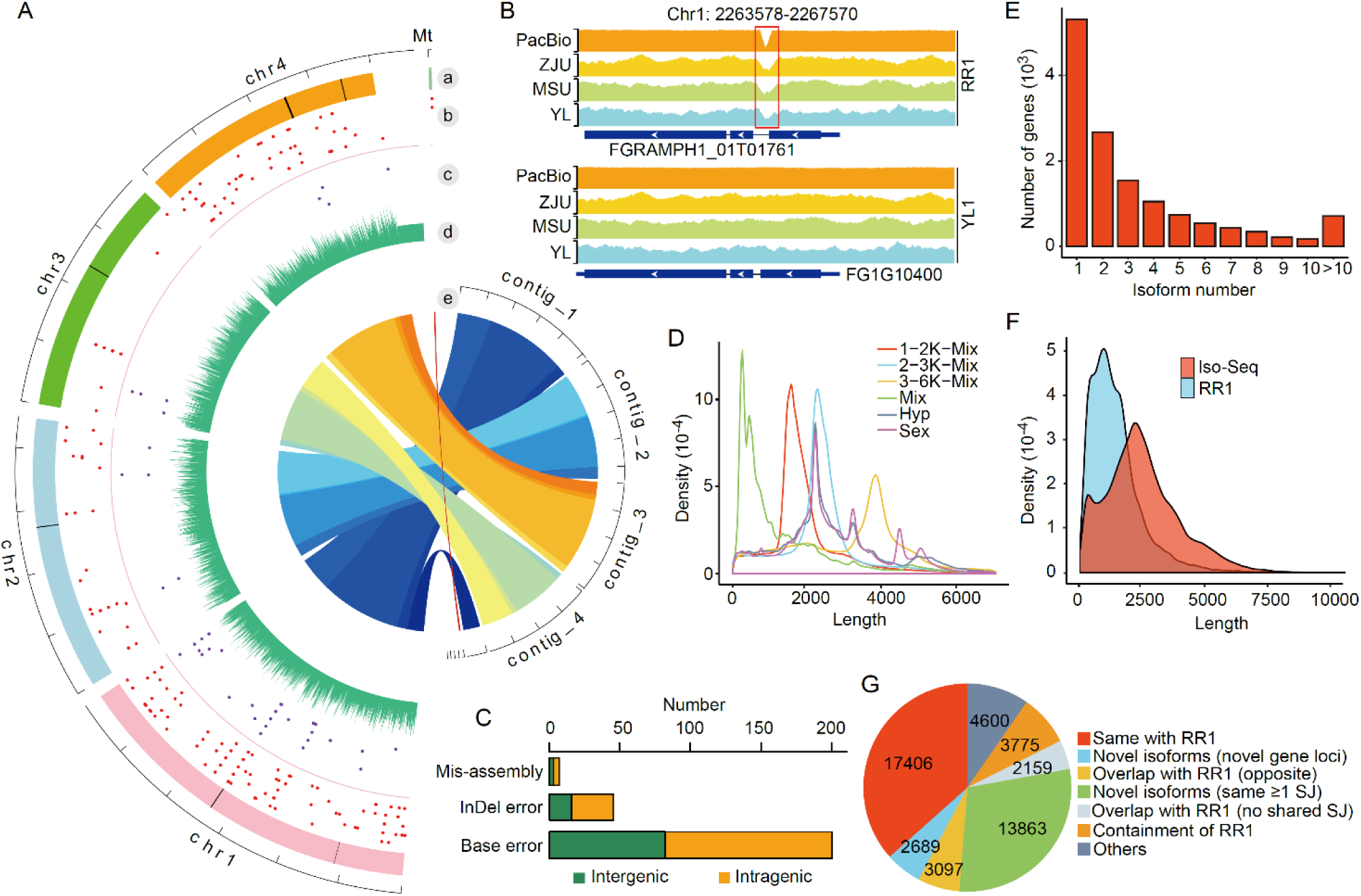
Improvement of the genome sequence of *F. graminearum* and its transcript annotation by PacBio SMRT sequencing. (*A*) Comparison of PacBio and RR1 assemblies of four chromosomes (chr1-chr4). a: Ideograms of RR1 chromosomes with mis-assembly regions indicated by black bars; b: Base error positions (red dots) in RR1 assembly; c: InDel error positions (purple dots) in RR1 assembly; d: PacBio read coverage in 10 kb bin; e: Syntenic blocks between PacBio and RR1 assemblies connected with ribbons. (*B*) An example of mis-assembly errors in the RR1 assembly. Coverage of Illumina reads from three PH-1 lab stocks (ZJU, MSU and YL) and PacBio reads from YL stock at the marked locus were mapped to the RR1 and YL1 assemblies. Red box highlights the mis-assembly region. (*C*) Number of corrected genomic errors located in the intragenic or intergenic regions. (*D*) Subread length distribution of different Iso-Seq libraries. (*E*) Distribution of the number of transcript isoforms per gene. (*F*) Comparison of transcript length distributions between the RR1 annotations and Iso-Seq transcript isoforms. (*G*) Classification of Iso-Seq transcript isoforms.

### Generating a comprehensive transcript annotation by Iso-Seq and strand-specific RNA-Seq

The most recent *F. graminearum* annotation based on PH-1 reference genome RR1 (Ensembl Fungi) contains 14,145 protein coding genes, with each of them being associated with one transcript derived from computational prediction (King et al. 2017). To identify as many transcripts as possible, we combined equal amounts of poly(A)^+^ RNA purified from six cell types of PH-1 for PacBio SMRT sequencing, including perithecia collected at 6 days postfertilization (dpf) (Sex6d), conidiating cultures with phialides and conidia harvested from 5-day-old CMC cultures (CMC5d), germlings from 12 h YEPD cultures (YEPD12h), aerial mycelia from 5-day-old PDA (PDA5d) and carrot agar (CA) (CA5d) cultures, and DON-producing hyphae sampled from 3-day-old TBI cultures supplemented with arginine (TBI3d). To increase the representation of different mRNA populations, three size-selected (1-2 kb, 2-3 kb, and 3-6 kb) and one non-size selected Iso-Seq libraries were constructed and sequenced on the PacBio Sequel System (Supplemental Table S1). Furthermore, to facilitate comparison between vegetative growth and sexual reproduction, we also performed non-size selected Iso-Seq with poly(A)^+^ RNA isolated from 24-h hyphae and 6-dpf perithecia with two independent biological replicates. In total, 5,423,051 reads of insert (ROIs) were yielded from these Iso-Seq libraries (Supplemental Table S1). The three size-selected libraries had the expected distribution of transcript lengths (Fig. 1D). To analyze the Iso-Seq data, we developed a computational pipeline (see Methods and Supplemental Fig. S1) and used it to obtain 47,589 high-quality unique transcript isoforms representing 13,712 genes.

Over 60% (8,391) of genes in the Iso-Seq transcript isoform set had two or more transcript isoforms with an average of 3.5 (Fig. 1E). Remarkably, one gene (FG1G24420) had more than 200 transcript isoforms. On average, the Iso-Seq transcript isoforms (median 2,341 nt) were 160% longer than the predicted RR1 transcripts (median 1,462 nt) (Fig. 1F), which is mainly due to increases in the length of UTRs because less than half of RR1 transcripts have complete 3’- and 5’-UTR annotations. We then compared and categorized the most closely matching RR1 transcripts using the *GffCompare* utility (Pertea and Pertea 2020). As a result, 36.6% (17,406) of the Iso-Seq transcript isoforms were same with the RR1 transcripts with respect to the intron chain (Fig. 1G). However, 63.4% (30,183) of the Iso-Seq transcript isoforms were not present in or different from RR1 transcripts. Among them, 5,786 isoforms (12.2%) represent novel genes either from novel loci (2,689) or overlap with existing gene loci on the opposite strand (3,097), and 16,022 isoforms (33.6%) are possible novel isoforms of existing genes that share at least one splice site with RR1 transcripts but differ at other splice sites (13,863) or overlap with the RR1 transcripts but with different splice sites (2,159). In addition, 3,775 Iso-Seq transcript isoforms (7.9%) were found to be longer than RR1 transcripts and have extra splice sites. Therefore, the high proportion of novel genes or isoforms identified in our Iso-Seq transcript isoform set suggests the presence of significant transcriptional diversity that has not been identified in previous studies in *F. graminearum*.

Totally, the Iso-Seq transcript isoform set covered 77.5% (10,958) of existing RR1 genes. The remaining 3,187 genes had no Iso-Seq transcripts. These genes were generally had lower expression levels compared to the genes with Iso-Seq transcripts (Supplemental Fig. S2). To obtain more complete transcript annotations, we independently generated strand-specific RNA-Seq data by Illumina for each of the poly(A)^+^ RNA samples of the six cell types (Supplemental Table S2). After transcript assembly of these RNA-Seq data, we generated 1,003 full-length transcripts for 882 RR1 genes that had no Iso-Seq transcripts, and 300 full-length transcripts from novel gene loci with reliable expression levels (≥5 TPM). In addition, for 232 Iso-Seq transcript isoforms that likely had incomplete 5’-end, the corresponding assembled full-length transcripts were obtained.

By combining the full-length transcripts from both Iso-Seq and strand-specific RNA-Seq together with the remanent 2,305 RR1 genes without full-length transcripts available, we generated a comprehensive reference annotation for *F. graminearum* PH-1 that has 51,617 transcript isoforms for 17,189 genes. To further annotate CDSs of protein coding genes, we used *getorf* from EMBOSS package (Rice et al. 2000) to predict open reading frame (ORF) for each transcript isoform. A total of 50,007 transcript isoforms representing 16,547 genes have predicted ORF of ≥30 aa, including 2,998 novel protein-coding genes discovered in this study. Among these, 285 had BLASTp hits and 5 had Pfam matches (E value<0.001) (Supplemental Table S4). The extensive update of the annotation was adopted in a new gene naming schema according to gene’s chromosome order and location (e.g., FG1G00010), and associated with legacy annotation gene IDs from Broad Institute V3 (FGSG_XXXXX) and Ensembl Fungi RR1 (FGRAMPH1_XXXXXXXX). In the following manuscript, we denoted this final set of annotation as the YL1 annotation. To facilitate its usage, we created the FgBase (fgbase.wheatscab.com), a *Fusarium graminearum* genome database that allows users to browse (JBrowse), search, and download the updated genome and annotation reported in this study.

### Landscape of alternative splicing

Based on the new YL1 reference annotation, we performed a systematic characterization of alternative splicing (AS) in *F. graminearum*. A total of 54,613 AS events were identified in the new YL1 annotation by *AStalavista* (Foissac and Sammeth 2007). These AS events affected 42.7% (4,997) of intron-containing genes and resulted in 17,229 splice isoforms. The numbers of AS events and transcript isoforms identified in this study are significantly higher than what have been previously reported in *F. graminearum* and other fungi (Zhao et al. 2013; Grutzmann et al. 2014), suggesting that the importance of AS in increasing fungal transcriptome complexity has been underestimated. Gene Ontology (GO) enrichment analysis showed that these AS genes are highly enriched for diverse biological processes, including glycolysis, chitin biosynthesis, inositol lipid-mediated signaling, potassium ion transmembrane transport, divalent metal ion transport, regulation of response to stimulus, TOR signaling, hexose metabolism, and positive regulation of transcription by RNA polymerase II (Supplemental Fig. S3).

We further classified the AS events into four basic types: intron retention (IR), alternative 5’-donor (A5), alternative 3’-acceptor (A3), and exon skipping (ES) (Kim et al. 2008) (Fig. 2E). Consistent with previous findings in fungi (Grutzmann et al. 2014), IR comprised the majority (61.9%) of AS events. A3 was the second most common (17.1%) AS events, followed by A5 (13.3%). The number of ES events was the least (7.7%). In addition, we found that the transcripts of 1,960 genes have multiple combinatory AS events, suggesting that more complex AS events also occurred frequently in *F. graminearum*. A total of 25,069 AS events were found in the sexual stage (perithecia) while near four times less AS events (6,552) were during vegetative growth (hyphae) (Fig. 2A). Interestingly, the relative proportion of IR events was higher in the vegetative growth stage.

**Figure 2.**
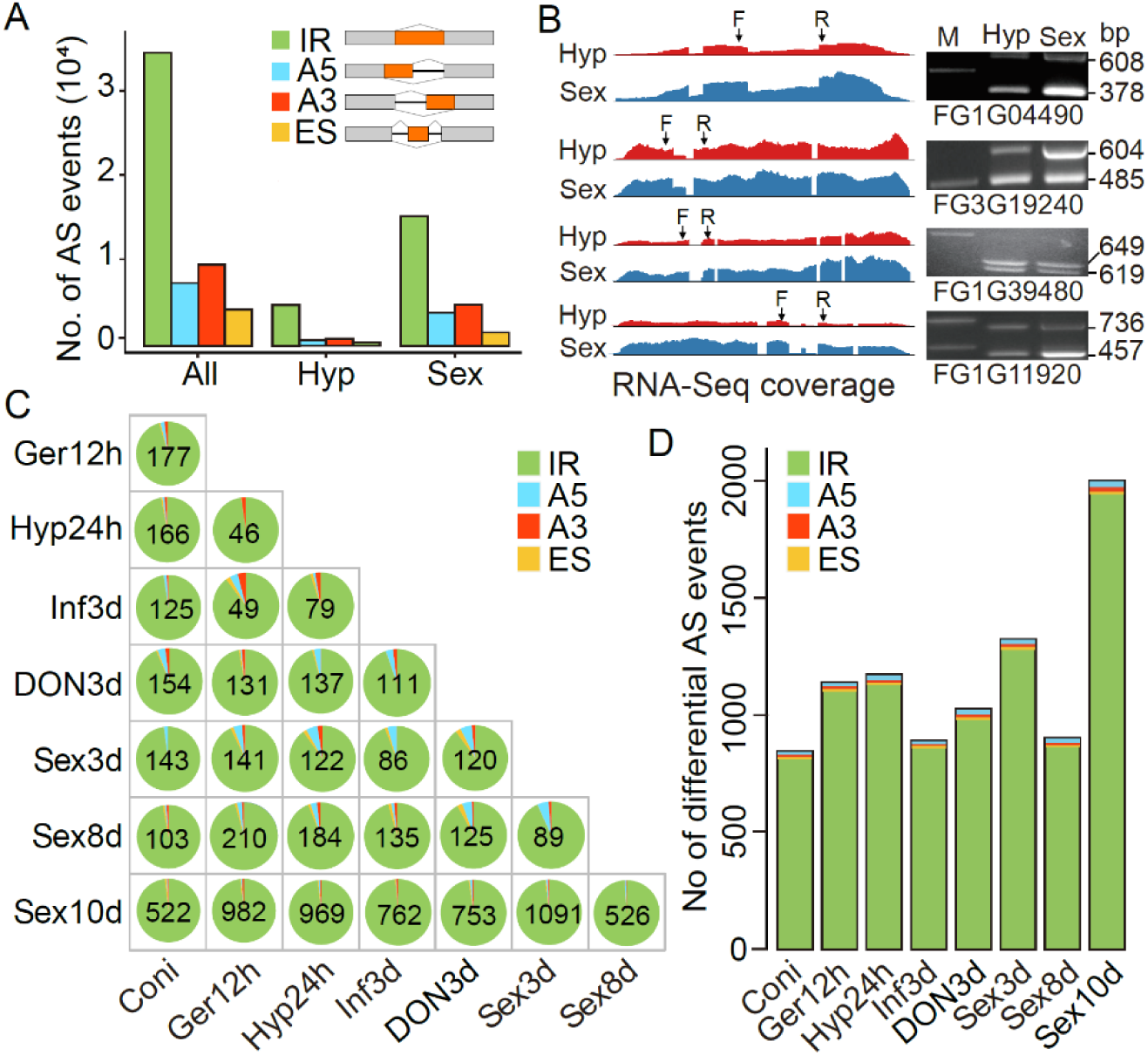
Landscape of alternative splicing (AS) in various cell types. (*A*) Distribution of four types of AS events, including intron retention (IR), alternative 5’-donor (A5), alternative 3’-acceptor (A3), and exon skipping (ES) in transcripts of all samples (All) as well as vegetative hyphae (Hyp) and sexual perithecia (Sex) samples. (*B*) RT-PCR validation of the four types of AS events in the marked loci with RNA isolated from hyphae (Hyp) and perithecia (Sex). The expected size for each band is indicated. M, DNA makers. RNA-Seq read coverage of the four associated genes is shown in left panel. PCR primers (F, forward and R, reverse) are designed to flank the splicing events. (*C*) Number of significantly differentially AS events between pairs of marked samples (FDR<0.05). Pie charts show the proportion of differential IR, A5, A3, and ES events. (*D*) Number of significantly non-redundant differential AS events relative to each of the marked samples (FDR<0.05). For *C*-*D*, see Supplemental Table S2 for details of sample information.

To validate the accuracy of the AS events detected with Iso-Seq reads, we randomly selected four examples that showed each type of AS events for reverse transcription (RT)-PCR and Sanger sequencing. Primers (Supplemental Table S5) suitable for distinguishing different splice isoforms were designed and used for RT-PCR with RNA from vegetative hyphae and perithecia. For all the samples, when the resulting PCR products were separated on gels, we observed bands of expected sizes based on the AS events identified in Iso-Seq data (Fig. 2B). All these four AS events selected for verification were further confirmed by Sanger sequencing analysis with the amplified fragments. Interestingly, we found that the relative brightness of the two bands from vegetative hyphae and perithecia were different for genes FG3G19240 and FG1G11920, indicating that expression of splice isoforms may exhibit a stage-preferential pattern.

When Iso-Seq full-length transcript isoforms from sexual stage were compared with those from vegetative growth stage, 12,602 transcript isoforms from 6,032 genes were found to be common between these two stages (Supplemental Fig. S4A). Nevertheless, A total of 14,080 transcript isoforms (52.8%) associated with 3,291 genes were stage-specific and identified only in perithecia or vegetative hyphae. Sexual stage had the higher proportion of stage-specific transcript isoforms (40.7%). Remarkably, one gene FG1G24420 was found to express 197 transcript isoforms during sexual reproduction but only one in vegetative hyphae. GO enrichment analysis showed that genes with sexual stage-specific transcript isoforms were significantly enriched for regulation of transcription, oxidation-reduction, and transmembrane transport processes, while genes with vegetative growth stage-specific transcript isoforms were significantly enriched for RNA processing, methylation, and RNA modification processes (Supplemental Fig. S4B).

### Differential AS events in different cell types

To examine the dynamic of AS events across different cell types, we calculated a Percent Spliced-In (PSI) index using *SUPPA2* (Trincado et al. 2018) for each AS event in different samples. PSI index calculated as the fraction of the inclusion reads to the total reads (both inclusion and exclusion reads) to measure the inclusion level of a given splicing event. Hierarchical clustering revealed that the PSI values were variable in different samples (Supplemental Fig. S4C), suggesting that the AS events are subject to cell type-specific controls.

To identify differential AS events in different cell types, we generated replicate strand-specific RNA-Seq data from conidia (Coni), 12 h conidial germlings (Ger12h), 24 h vegetative hyphae (Hyp24h), infection hyphae 3 days post inoculation (dpi) (Inf3d), 3-day-old DON production hyphae in TBI culture supplemented with NH_4_NO_3_ (DON3d), and three sexual stages (Sex3d, Sex8d, and Sex10d) (Supplemental Table S2). We then used *CASH* (Wu et al. 2018) to detect differential AS events among these samples based on the YL1 annotation. Pairwise comparative analysis revealed that the number of differential AS events ranged from 46 to 1,091 among these samples (FDR<0.05), with the largest between Sex3d and Sex10d comparison and the least between Ger12h and Hyp24h (Fig. 2C). Totally, the non-redundant differential AS events relative to each sample ranged from 844 to 1,996 (Fig. 2D). The Sex10d sample had the greatest number of differential AS events in comparison with other samples, suggesting a distinct AS landscape in the late stage of sexual reproduction.

### A global increase in intron inclusion in aging or dormant tissues

The vast majority (>95%) of differential AS events are IR events (Fig. 2D), indicating that IR is more likely to be regulated across different cell types than other types of AS. The differential IR events can be further subdivided into two opposite types: intron inclusion (increased intron retention) and intron exclusion (decreased intron retention). Interestingly, in Ger12h, Hyp24h and Inf3d, no more than 12% of the differential IR events was intron inclusion whereas the fraction of intron inclusion was over 33% in Coni, Sex8d, and Sex10d (Fig. 3A). During sexual reproduction, intron inclusion continuously increased from Sex3d to Sex10d. In Sex10d, 95% of the differential IR events were intron inclusion (Fig. 3A). Consistent with these observations, hierarchical clustering of PSI values from each sample revealed that the PSI values of most AS events were higher in Sex10d but lower in Ger12h, Hyp24h and Inf3d (Fig. 3B). In comparison with Ger12h, Hyp24h and Inf3d that have active growth, Sex10d and Coni were representative of aging and dormant states (Wang et al. 2021a). Therefore, these observations suggest that increasing abundance of intron-retained isoforms is associated with aging or dormant states.

**Figure 3.**
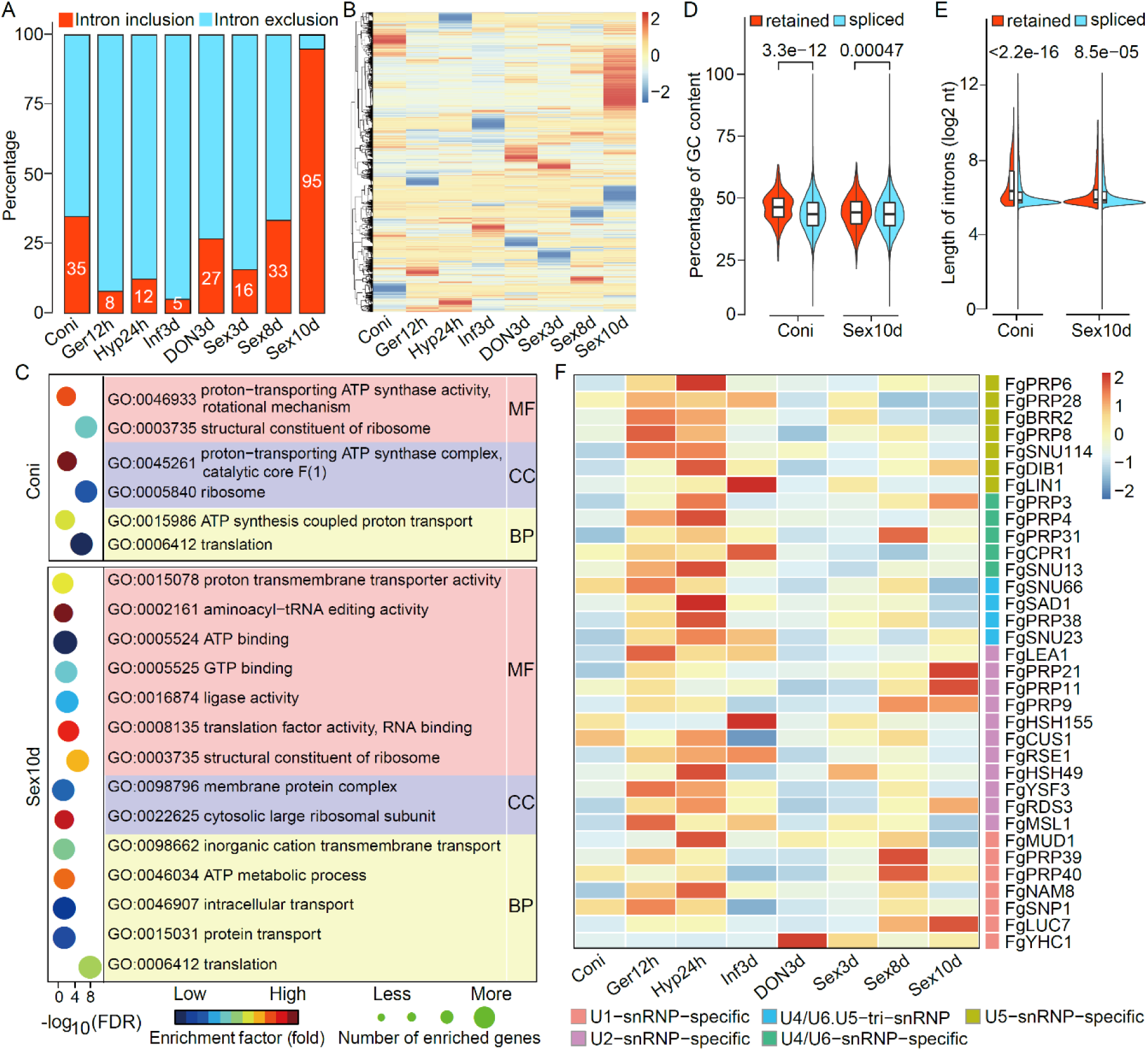
Regulation of intron retention in *F. graminearum*. (*A*) Proportion of intron inclusion and intron exclusion in the non-redundant differential intron retention events relative to each of the marked samples. (*B*) Heatmap of Percent Spliced-In (PSI) values across different samples. High (orange to red) and low (yellow to blue) PSI values are depicted as Z-scores for each AS event. (*C*) Enriched GO terms in genes with differentially retained introns in conidia (Coni) or 10-dpf perithecia (Sex10d). MF, Molecular Function; CC, Cell Component; BP, Biological Process. (*D-E*) Comparison of GC contents (*D*) and intron lengths (*E*) between the spliced introns and differentially retained introns in conidia and 10-dpf perithecia. *P* values are from two-tailed Wilcoxon rank sum test. (*F*) Heatmap of the expression of putative genes encoding spliceosomal components in different samples. High (orange to red) and low (yellow to blue) expression levels are depicted as Z-scores for each gene. Subunits of the five canonical snRNP subcomplexes are indicated with different colors on the right. For *A, B*, and *D-F*, see Supplemental Table S2 for details of sample information.

Genes with increased intron inclusion in Coni were enriched for functions associated with ATP synthesis coupled proton transport and translation, while those in Sex10d were enriched for functions associated with ATP metabolic process, translation, inorganic cation transmembrane transport, intracellular transport, and protein transport (Fig. 3C). Furthermore, analysis of the retained introns increased in Coni and Sex10d revealed that they generally are longer and have higher GC contents compared to the spliced introns (Fig. 3D-E). Taken together, our results suggest that a global increase in intron inclusion is likely a transcriptional signature of aging or dormant states in *F. graminearum*. Interestingly, for most of the genes encoding spliceosomal components their expression levels were generally higher in the active tissues (Ger12h, Hyp24h and Inf3d) but lower in the old or quiescent tissues (Coni and Sex10d) (Fig. 3F). It is possible that depressed expression of spliceosomal genes is directly related to the increased intron retention in the old or dormant tissues.

### Majority of AS events alter ORFs in *F. graminearum*

We next assayed the alteration of encoding proteins in the AS transcript isoforms with respect to the canonical transcript isoform of each gene, which was defined as the transcript isoform with the highest expression levels in most samples. Based on the changes in the ORFs caused by AS events, the 12,232 AS transcript isoforms were divided into ten categories (Fig. 4A). The most frequent category is 5’-ORF shortening, which accounts for 26.4% of the AS transcript isoforms. In fact, 44.6% of AS transcript isoforms contain shortened ORFs at the 5’ and/or 3’ end. In contrast, only a minor proportion (4.4%) have lengthened ORFs at 5’ and/or 3’ end (Fig. 4A). Additionally, 11.6% of AS transcript isoforms have in-frame ORF alteration with the number of nucleotides lost or gained by AS events that can be divided by 3. Overall, the majority of AS transcript isoforms encoded altered ORF sequences relative to canonical transcript isoforms; only 21.5% shared the same ORF sequences as the canonical transcript isoforms.

**Figure 4.**
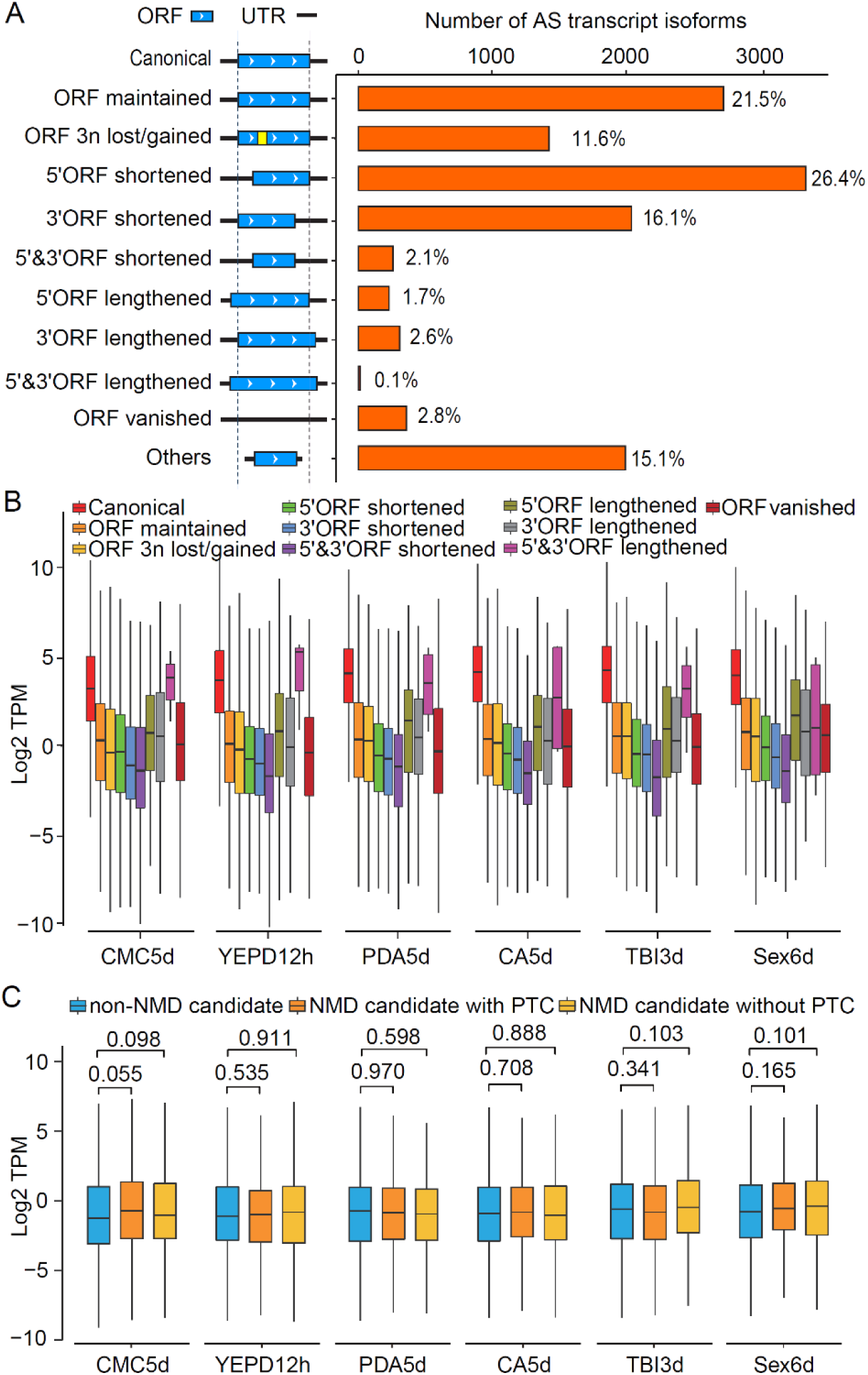
Alteration of encoding proteins and expression of alternative splicing (AS) transcript isoforms. (*A*) The number and percentage of AS transcript isoforms for each marked category. The diagrams in the middle shows the effects of each category of AS transcript isoforms on protein coding relative to the canonical transcript isoform. ORF, open reading frame; UTR, untranslated region. (*B*) Box plots comparing the expression levels (Log_2_ TPM) of different categories of AS transcript isoforms. (*C*) Box plots comparing the expression levels (Log_2_ TPM) of the predicted premature stop codon (PTC)-containing and non-PTC-containing NMD candidates and the non-NMD candidates of the 3’-ORF shortened AS transcript isoforms. *P* values are from two-tailed Kruskal-Wallis test. For *B-C*, see Supplemental Table S2 for details of sample information.

We then examined the expression levels of transcript isoforms belonging to different categories and found that the AS transcript isoforms with shortened ORFs commonly had lower expression levels in comparison with other categories of transcript isoforms. In comparison with other isoforms, the 5’or 3’-ORF shortened transcript isoforms had the lowest expression levels in all the samples (Fig. 4B), implying that these categories of AS transcript isoforms may be subjected to degradation by RNA surveillance mechanisms.

### AS may be not coupled to NMD in *F. graminearum*

In addition to increasing proteomic diversity, AS may also regulate gene expression by generating transcript isoforms that are recognized and degraded by mRNA surveillance mechanisms. One of these mechanisms is nonsense-mediated mRNA decay (NMD) (Kishor et al. 2019), selectively degrading premature stop codon (PTC)-containing transcripts to prevent production of truncated proteins. In animals, transcripts with a PTC or even a normal stop codon >50 nt upstream of the last exon-exon junction can efficiently trigger NMD (Garcia-Moreno and Romao 2020). Using this rule, we identified 2,057 candidate NMD targets from the AS transcript isoforms. Among them, 764 (37.1%) contained a PTC, accounting for 6.2% of the total AS transcript isoforms. Surprisingly, the expression levels of both PTC-containing and non-PTC-containing NMD candidates were not significantly lower than that of non-NMD candidates in the 3’-ORF shortened transcript isoforms (Fig. 4C), indicating that the AS transcript isoforms with NMD-eliciting features may be not recognized and degraded by the NMD pathway in *F. graminearum*.

To further characterize the NMD in *F. graminearum*, we identified and deleted *FgUPF1* (FG1G42730) (Fig. 5A; Supplemental Table S6 and S7), an ortholog of the yeast *UPF1* gene encoding the central factor of the NMD pathway (Kishor et al. 2019). The Δ*Fgupf1* mutant grew much slower than the wild type on PDA plates (Fig. 5B), demonstrating the importance of *FgUPF1* in *F. graminearum*. We then analyzed expression of the predicted NMD targets in hyphal samples of the wild type and Δ*Fgupf1* mutant using strand-specific RNA-Seq (Supplemental Table S2). The expression ratios of the PTC-containing NMD targets (TPM of NMD-target/total TPMs of all transcripts per gene) were not increased in the Δ*Fgupf1* mutant compared to the wild type (Fig. 5C). A total of 734 transcripts were up-regulated over 2-fold in the Δ*Fgupf1* mutant (|log_2_ fold change|≥1, q-value<0.05) (Fig. 5D). The vast majority of these transcripts were canonical transcript isoforms or AS transcript isoforms without ORF changes (Fig. 5E). The 3’-ORF shortened transcript isoforms, however, were rarely affected by deletion of *FgUPF1*. In addition, transcripts with upstream ORF (uORF) or long 3’-UTR were also reported to induce NMD (Garcia-Moreno and Romao 2020). We therefore analyzed the occurrence of uORFs in the transcripts and their 3’-UTR lengths. There was no obvious difference for the proportion of uORF-containing transcripts in the transcripts up-regulated in the Δ*Fgupf1* mutant compared to non-differentially expressed transcripts (Fig. 5F). In general, the 3’-UTR sequences of the transcripts up- or down-regulated in the Δ*Fgupf1* mutant were shorter than those of non-differentially expressed transcripts (Fig. 5F). These results indicate that AS may be not coupled to NMD in *F. graminearum*.

**Figure 5.**
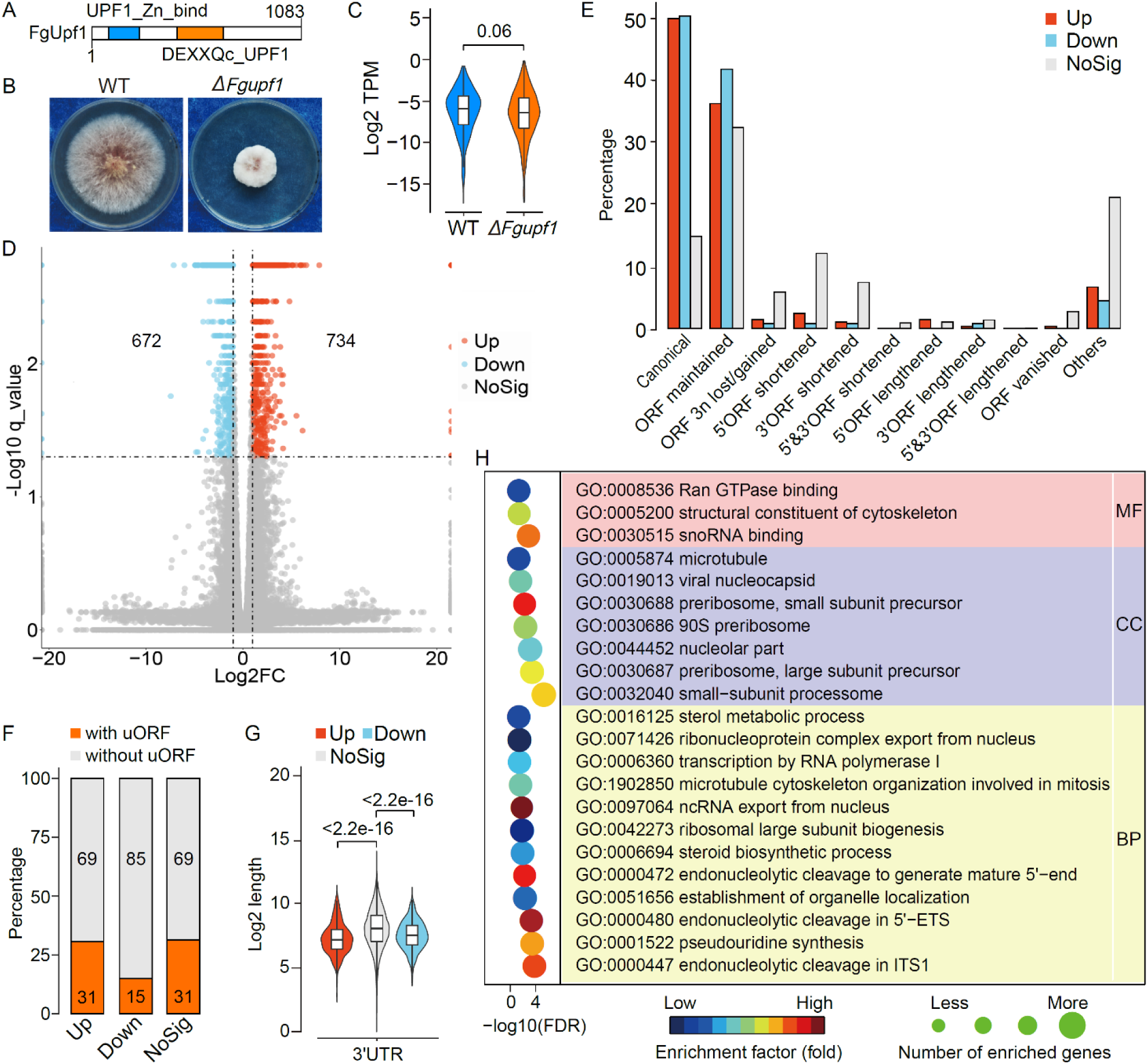
Characterization of the Δ*Fgupf1* mutant. (*A*) Conserved domains of FgUpf1. DEXXQc_UPF1, DEXXQ-box helicase domain of Upf1; UPF1_Zn_bind, RNA helicase (Upf2 interacting domain). (*B*) Three-day-old cultures of the wild-type strain PH-1 (WT) and Δ*Fgupf1* mutant grown on potato dextrose agar (PDA) plates. (*C*) Box plots comparing the expression levels (Log_2_ TPM) of the predicted PTC-containing NMD candidates in WT and Δ*Fgupf1* mutant. *P* value is from two-tailed Wilcoxon rank sum test. (*D*) Volcano plot of the significantly up- and down-regulated transcripts in the Δ*Fgupf1* mutant (|log_2_ fold change| ≥1, q-value<0.05). (*E*) Distribution of the up-regulated, down-regulated, and non-differentially expressed (NoSig) transcripts in the Δ*Fgupf1* mutant in each marked category. (*F-G*) Percentage of transcripts with upstream ORF (uORF) (*F*) and box plots comparing the 3’-UTR lengths (*G*) in the same three categories of transcripts (Up, Down, and NoSig). *P* values are from two-tailed Kruskal-Wallis test. (*H*) Enriched GO terms in genes with down-regulated transcripts in the Δ*Fgupf1* mutant. MF, Molecular Function; CC, Cell Component; BP, Biological Process.

Interestingly, GO enrichment analysis revealed that the 672 transcripts down-regulated in Δ*Fgupf1* mutant were mostly enriched for cell components associated with ribosome biogenesis, including GO terms of “90S preribosome”, “small-subunit processome”, “preribosome, small subunit precursor”, “preribosome, large subunit precursor”, and “nucleolar part” (Fig. 5G). Therefore, it is likely that *FgUPF1* regulates the expression of genes related to ribosome biogenesis in *F. graminearum*, which may be directly related to the severe growth defect of the Δ*Fgupf1* mutant.

### Landscape of alternative polyadenylation

Alternative polyadenylation (APA) can enhance transcriptome complexity by generating RNA isoforms that differ in their 3’ end. Since the polyadenylation sites (PASs) of mRNAs are well represented in the full-length, non-artificial-concatemer (FLNC) reads, we characterized the global polyadenylation events in the *F. graminearum* genome. Totally, 364,513 unique PASs were identified (Fig. 6A). Among these, 83.9% are in the 3’ UTR regions, resulting in variations in the length of 3’ UTRs. About 16% of the unique PASs are in the CDS (14.2%) and 5’ UTR (1.4%) regions. Such APA isoforms may produce truncated proteins or had no protein products. Nevertheless, among the FLNC reads, only 4.1% of them had non-3’ UTR PASs (Fig. 6A), suggesting that the vast majority of APA events affect only the length of 3’ UTRs in *F. graminearum*.

**Figure 6.**
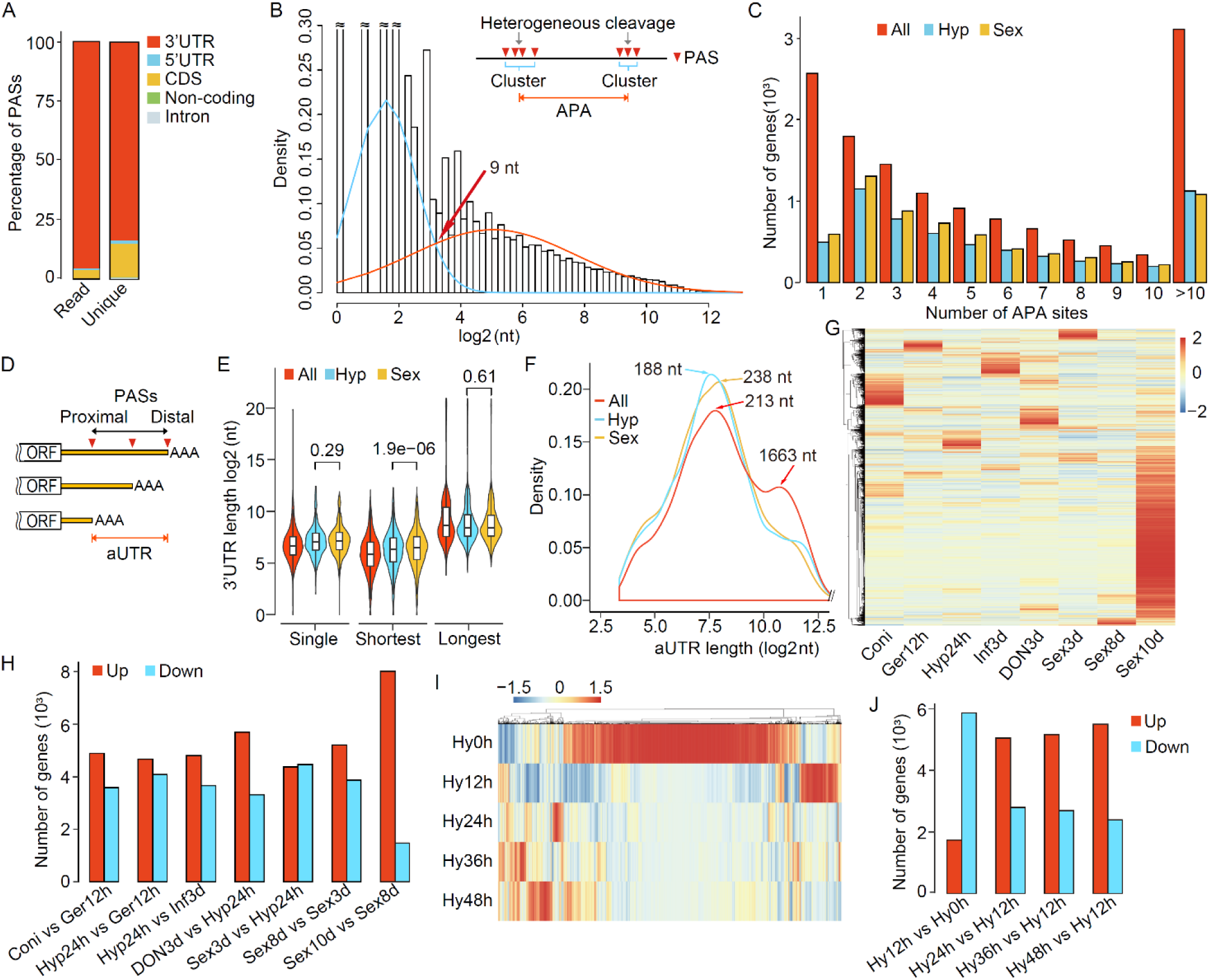
Landscape and regulation of alternative polyadenylation (APA) in *F. graminearum*. (*A*) Statistics of FLNC reads mapped to different regions of the *F. graminearum* genome and unique genomic polyadenylation sites (PASs). (*B*) Distribution of distances between adjacent PASs, with the blue line showing the distance between PASs within a PAS cluster and the red line showing the distance between PASs from different PAS clusters. The red arrow marks the crossover point used to group PASs into a PAS cluster. The PASs (arrowheads) within a PAS cluster were considered to be derived from heterogeneous cleavage while the PASs between different PAS clusters were considered to be derived from APA. (*C*) Distribution of the number of APA sites per gene detected in all samples (All) as well as in vegetative hyphae (Hyp) and sexual perithecia (Sex). (*D*) Schematic drawing of three APA isoforms derived from different PASs (arrowheads) in the 3’-UTR. The region between the first (most proximal) PAS and last (most distal) PAS is named alternative 3’-UTR (aUTR). (*E*) Box plots comparing the 3’-UTR length of genes without APA (Single) and genes with APA (only the shortest and longest 3’-UTR isoforms were plotted). (*F*) Distribution of the length of aUTRs identified in all samples (All) as well as in vegetative hyphae (Hyp) and sexual perithecia (Sex). The peak values are indicated. (*G*) Heatmap of the M/m values for each PAS across marked samples. High (orange to red) and low (yellow to blue) M/m values are depicted as Z-scores for each PAS. (*H*) Number of genes with significantly increased (Up) or decreased (Down) distal PAS usage in marked pairwise comparisons of different samples (*p-*value<0.05). (*I*) Heatmap of the M/m values for each PAS in conidia (Hy0h) and vegetative growth from germlings (Hy12h) to older hyphae (Hy48h). High (orange to red) and low (yellow to blue) M/m values are depicted as Z-scores for each PAS. (*J*) Number of genes with significantly increased (Up) or decreased (Down) distal PAS usage in marked pairwise comparisons (*p-*value<0.05). For *G-J*, see Supplemental Table S2 for details of sample information.

Because heterogeneous cleavage by mRNA 3’-end processing machinery often leaves multiple PASs located close to one another, we thus modeled two types of distances between adjacent PASs and identified 9 nucleotides (nt) as the cutoff to distinguish APA from heterogeneous cleavage (Fig. 6B). Nearby PASs within 9 nt of each other in the same gene were clustered and defined as a PAS cluster (heterogeneous cleavage). In total, we identified 104,813 PAS clusters with an average of 6.1 clusters per gene. In each cluster, the PAS with most supporting FLNC reads was selected for subsequent analysis. In total, 11,133 genes (64.8%) displayed APA, including 6,123 in sexual stage and 5,530 in vegetative hyphae (Fig. 6C). On average, there are 9.2 PAS clusters per gene (Fig. 6C), indicating extensive APA events in *F. graminearum* during both sexual reproduction and vegetative growth.

APA sites in 3’-UTRs lead to transcript isoforms with different 3’-UTR lengths. The most proximal (first) and most distal (last) PASs generate shortest and longest 3’-UTRs, respectively (Fig. 6D). For the APA genes in *F. graminearum*, the median length of the shortest and longest 3’ UTRs were 58 and 395 nt, respectively (Fig. 6E). Compared to vegetative hyphae, the shortest 3’-UTR isoforms were longer in sexual stage (a median of 90 vs 81) but the longest ones had no significant difference (a median of 344 vs 345). Furthermore, the distance between the first and last PASs in 3’ UTR was named alternative 3’-UTR (aUTR) (Fig. 6D). The aUTRs in sexual stage (peak value, 238 nt) were 127% longer than those in vegetative growth stage (peak value, 188 nt) (Fig. 6F). Interestingly, the distribution of aUTR lengths for all samples had two peaks: the main one at 213 nt and the minor one at 1,663 nt (Fig. 6F). It is likely that unusual long aUTRs occurred in certain cell types except perithecia and vegetative hyphae.

### Increased expression ratio of long 3’ UTR isoforms in aging or dormant tissues

The relative abundance of different APA isoforms of each gene was analyzed with *roar* (Grassi et al. 2016). The M/m value was calculated as the expression ratio of the distal PAS isoform (M) to the proximal PAS isoform (m) for each PAS except the last PASs (The larger the M/m value, the higher abundance for the distal PAS isoform). For majority of PASs, the M/m value was less than 1 (Supplemental Fig. S5), suggesting that the proximal PAS isoforms (with shorter 3’ UTR) are generally more abundant than the distal PAS isoforms (with longer 3’ UTR) in *F. graminearum*.

Notably, the median of M/m values in Sex10d were obviously larger than that in other samples (Supplemental Fig. S5). Consistent with these observations, hierarchical clustering revealed that most of the M/m values were largest in Sex10d (Fig. 6G), suggesting that the usage of distal PASs is generally increased in Sex10d. Furthermore, we identified the genes with significantly differential PAS usage among different samples, and found that genes with increased distal PAS usage in the older or dormant tissues as compared to younger tissues outnumbered those with decreased distal PAS usage (Fig. 6H). To further support these observations, we generated additional strand-specific RNA-Seq data with RNA samples isolated from conidia (0 h), germlings (12 h), and gradually aging hyphae (24 h, 36 h, and 48 h) (Supplemental Table S2). In comparison with 12 h germlings, the usage of distal PASs was remarkably increased in dormant conidia and also continuously increased in aging hyphae from 24 h to 48 h (Fig. 6I-J). These results suggest that increasing expression of long 3’-UTR isoforms is associated with aging or dormant state.

Genes with increased distal PAS usage in Sex10d, Coni, and older hyphae 48 h were then analyzed for enriched GO terms. A great number of GO terms related to autophagy, signal transduction, RNA binding, RNA processing, RNA splicing, RNA metabolism, translation, protein folding, protein localization, protein transport, protein binding, protein phosphorylation, ubiquitin-mediated proteolysis, chromatin organization, and histone modification were found to be significantly enriched in the genes from Sex10d, Coni, and/or hyphae 48 h (Supplemental Table S8). Many of these functions or pathways have been reported to be associated with cellular senescence or aging in animals (Deschenes and Chabot 2017; Chen et al. 2018; Angarola and Anczukow 2021). Therefore, it is possible that 3’-UTR lengthening may act as a novel mechanism in regulating aging and dormancy in *F. graminearum*.

### Polyadenylation signals surrounding PASs

To search for *cis*-acting elements that may guide cleavage and polyadenylation, we extracted upstream 100 nt and downstream 100 nt sequence surrounding each PAS to examine nucleotide distributions and enriched motifs. Apparently, the single-nucleotide frequencies in *F. graminearum* are similar to that observed in yeasts (Vavasseur and Shi 2014; Liu et al. 2017) (Fig. 7A). These included a U/A-rich region (efficiency element) within 75 nt upstream of the PAS, an A-rich peak (positioning element) located -25 to -10 nt, a U-rich peak (upstream U-rich element) immediately before the PAS, a short A-rich peak from +2 to +5 nt, and a U-rich region (downstream U-rich element) after the A-rich peak. Four enriched hexameric motifs, AATWVA (W = A or T, V = A, C or G), TAKMTA (K = G or T, M =A or C), TTTTTT, and HGTGAH (H = A, C or T) (Fig. 7B) were found to peak in the positioning, efficiency, upstream U-rich, and downstream U-rich elements in *F. graminearum*, respectively (Fig. 7C). The AATWVA and TTTTTT motifs occurred in a highly position-specific manner, suggesting that they are mechanistically important for cleavage and polyadenylation in *F. graminearum*.

**Figure 7.**
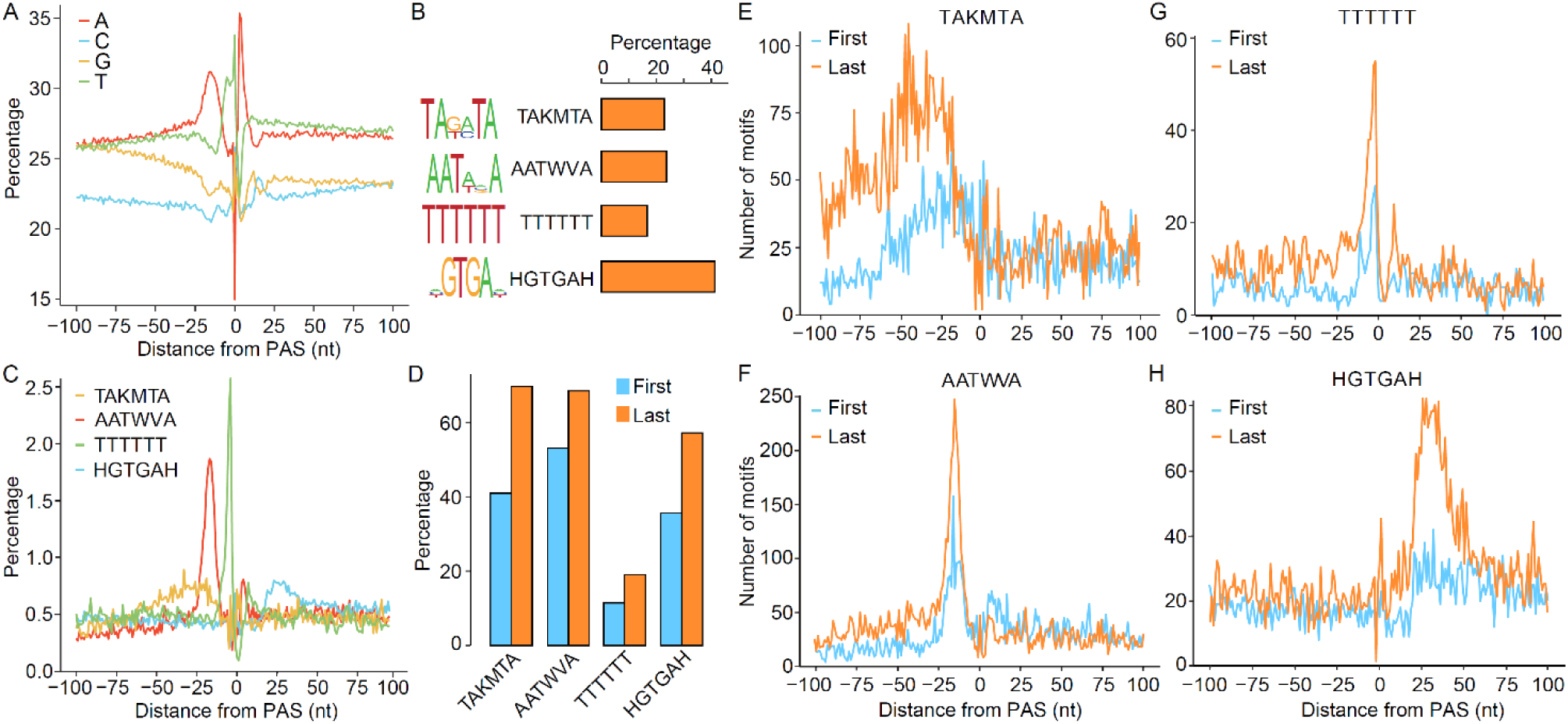
*Cis*-acting elements surrounding the polyadenylation sites (PASs). (*A*) Nucleotide frequencies around the PASs. (*B*) Four enriched hexameric motifs and percentage of PASs with these motifs. (*C*) Distribution of four hexameric motifs in the flanking region of PASs. (*D*) Comparison of the percentage of the four hexameric motifs detected in the vicinity of the first and last PASs that are located in the same terminal exon. (*E-H*) Comparison of the distribution of each hexameric motif in the vicinity of the first and last PASs in the same terminal exon.

We then examined whether proximal and distal PASs in *F. graminearum* APA genes were surrounded with different polyadenylation signals by comparing the first and last PASs that are located in the same terminal exon for their surrounding motifs. Whereas no additional motifs were detected beyond the four motifs aforementioned, their occurrence was more abundant at distal PASs (Fig. 7D-H), indicating that distal PASs are stronger than proximal ones in *F. graminearum*.

### Regulatory roles of 3’-end processing factors in PAS selection

Upon *cis*-acting elements surrounding the PASs, the 3’-end processing machinery also plays important roles in APA regulation by influencing PAS selections. This machinery includes three major complexes that are necessary for cleavage and polyadenylation in yeast: Cleavage Factor IA (CFIA), Cleavage Factor IB (CFIB), and Cleavage and Polyadenylation Factor (CPF) (Vavasseur and Shi 2014). We examined the expression of the orthologs of the core 3’-end processing factors from these complexes in *F. graminearum*. These genes were found to have higher expression levels in the younger tissues but lower expression levels in the older or dormant tissues (Fig. 8A), suggesting that increased distal PAS usage in aging and dormant tissues may be due to global down-regulation of these core 3’-end processing factors.

**Figure 8.**
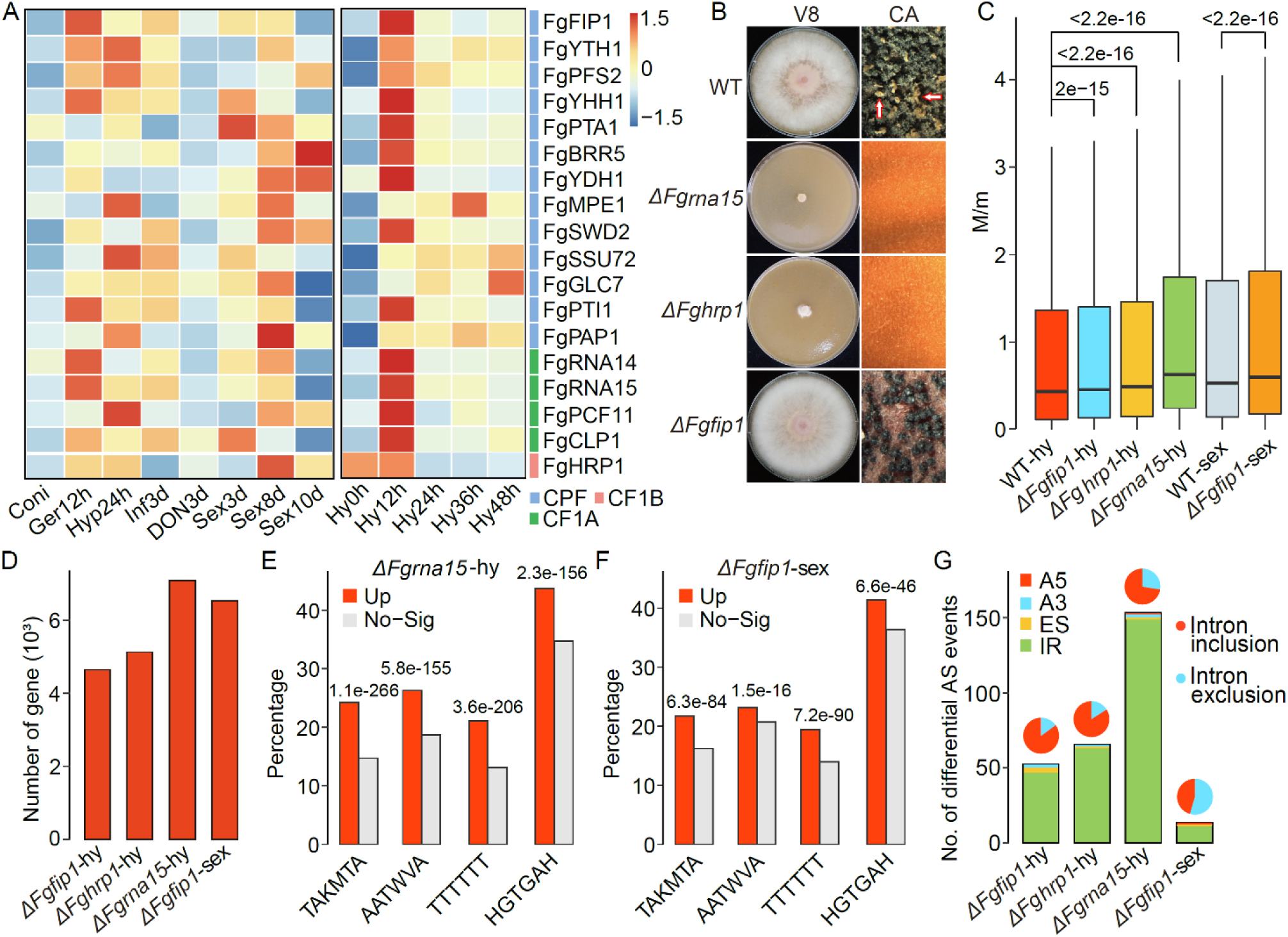
Roles of 3’-end processing factors in PAS selection and intron splicing. (*A*) Heatmap of the expression of genes encoding core mRNA 3’-end processing factors in different samples. High (orange to red) and low (yellow to blue) expression levels are depicted as Z-scores for each gene. Subunits of three complexes (CPF, CP1B, and CF1A) are indicated with different colors as marked. (*B*) Defects of the Δ*Fgfip1*, Δ*Fghrp1*, and Δ*Fgrna15* mutants in growth and sexual reproduction. Three-day-old V8 cultures and perithecia formed on carrot agar (CA) plates at 8 d post-fertilization. Arrows point to ascospore cirrhi. (*C*) Box plots of the M/m values of PASs for the marked samples. *P* values are from two-tailed Kruskal-Wallis test. (*D*) Number of genes with significantly increased (Up) distal PAS usage in marked mutants relative to the wild type (*p-* value<0.05). (*E-F*) Percentage of the four hexameric motifs detected in the vicinity of the PASs with increased (Up) or non-differential (No-Sig) distal PAS usage in Δ*Fgrna15* mutant (*E*) and the sexual sample of the Δ*Fgfip1* mutant (*F*). *P* values are from Fisher’s exact test. (*G*) Number of significantly differential AS events in marked mutants relative to the wild type (FDR<0.05). Pie charts show the proportions of differential intron inclusion and intron exclusion events. For *A, C, D*, and *G*, see Supplemental Table S2 for details of sample information.

To further characterize the function of the 3’-end processing machinery in APA regulation in *F. graminearum*, we selected to delete five of them including *FgRNA15, FgCLP1, FgHRP1, FgYTH1*, and *FgFIP1* (Supplemental Fig. S6), which are ortholog of *RNA15, CLP1, HRP1, YTH1*, and *FIP1* in *S. cerevisiae* (Preker et al. 1995; Barabino et al. 1997; Minvielle-Sebastia et al. 1997; Gross and Moore 2001; Pérez-Cañadillas 2006), respectively. Although all the five genes are essential for viability in *S. cerevisiae*, we obtained deletion mutants for the *FgRNA15, FgHRP1*, and *FgFIP1* genes (Supplemental Table S7). The *FgCLP1* and *FgYTH1* genes likely are also essential for viability in *F. graminearum* because we failed to identify deletion mutants after repeated attempts. The Δ*Fghrp1* and Δ*Fgrna15* mutants had severe growth defects and rarely produced aerial hyphae on V8 juice plates (Fig. 8B). They also failed to produce perithecia on mating plates, confirming their important roles in mRNA 3’-end processing in *F. graminearum*. Unexpectedly, the Δ*Fgfip1* mutant had no obvious defects in growth (Fig. 8B). Perithecia formed by the mutant were also normal in size and morphology but defective in ascospore release.

To investigate their roles in mRNA 3’-end processing, we performed strand-specific RNA-Seq analyses for hyphal sample of the Δ*Fgfip1*, Δ*Fghrp1* and Δ*Fgrna15* mutants and perithecium sample of the Δ*Fgfip1* mutant. In the resulting RNA-seq data, the median of M/m values was significantly increased in all three mutants compared to the wild type (Fig. 8C). We detected more than 4,000 genes with significantly increased distal PAS usage (increased M/m values) in each mutant (Fisher’s exact test, *p-*value<0.05) (Fig. 8D). These results suggest that the *FgRNA15, FgHRP1*, and *FgFIP1* genes are all functional in promoting proximal PAS usage. Especially, the median of M/m values and the number of genes with increased distal PAS usage were highest in the Δ*Fgrna15* mutant (Fig. 8C). Hierarchical clustering analysis also revealed that the M/m values of most PASs were dramatically increased in the Δ*Fgrna15* mutant (Supplemental Fig. S7), indicating the importance of *FgRNA15* in the regulation of the proximal PAS usage. Consistent with its obvious defect in sexual reproduction, the Δ*Fgfip1* mutant had a greater number of genes with increased distal PAS usage during sexual reproduction than in vegetative hyphae (Fig. 8D). Analysis of the motif occurrence revealed that the four motifs were significantly enriched in the vicinity of the PASs with increased M/m values in Δ*Fgrna15* mutant and the sexual sample of the Δ*Fgfip1* mutant (Fig. 8E-F). Similar situation was also observed in the hyphal sample of the Δ*Fgfip1* and Δ*Fghrp1* mutants except for the AATWVA motif, which was significantly depleted in the vicinity of the PASs with increased M/m values (Supplemental Fig. S8). These results suggest that all three genes are required for the recognition of these motifs in *F. graminearum*.

### *FgRNA15, FgHRP1*, and *FgFIP1* may regulate intron splicing in *F. graminearum*

During our analysis, we observed cases of intron splicing changed in the Δ*Fgrna15* mutant (Supplemental Fig. S9). We therefore performed transcriptome-wide identification of the significantly differential AS events in the three mutants in comparison with wild type. Totally, we detected 153, 65, and 52 differential AS events (FDR<0.05) in the hyphal sample of the Δ*Fgrna15*, Δ*Fghrp1*, and Δ*Fgfip1* mutants (Fig. 8G). Only 13 differential AS events were detected in the sexual sample of the Δ*Fgfip1* mutant. In all samples, the vast majority of detected differential AS events were IR events, and the majority of differential IR events in the hyphal sample belonged to the type of intron inclusion. These results suggest that the *FgRNA15, FgHRP1*, and *FgFIP1* genes may positively regulate intron splicing in *F. graminearum*. Especially, *FgRNA15* may play an important role in promoting intron splicing.

### Identification and characterization of lncRNAs in *F. graminearum*

In addition to protein-coding RNAs, non-coding RNAs constitute a major part of the transcriptome. In *F. graminearum*, only 547 long non-coding RNAs (lncRNAs) were identified by Illumina short-read sequencing (Kim et al. 2018). In this study, we identified 5,481 high-confidence lncRNAs from the Iso-Seq data (Fig. 9A), including 178 corresponding to previously discovered lncRNAs (Kim et al. 2018) (Fig. 9B). LncRNAs had a smaller number of exons compared to non-lncRNAs, with 77.4% (4,243) of them with a single exon (Fig. 9C). These lncRNAs were classified into four categories based on their positions: 40.3% (2,207) from intergenic regions, 14.9% (820) from the sense strand, 44.6% (2,445) from the antisense strand, and 0.2% (9) from intronic regions (Fig. 9A). A total of 819 lncRNAs were identified in sexual stage, near two folds more than 427 lncRNAs identified in the vegetative growth stage. In comparison with other stages, fewer intergenic but more genic (sense and antisense) lncRNAs were found in both vegetative growth and sexual stages. In general, the intergenic lncRNAs are shorter than genic lncRNAs, and the antisense lncRNAs tend to be longer than the sense ones (Fig. 9D). The lncRNAs in vegetative growth and sexual stages is generally longer than lncRNAs identified from other tissues. Additionally, the lncRNAs of sexual stage had relatively higher expression levels than those of vegetative growth stage (Fig. 9E). Hierarchical clustering revealed that the lncRNAs exhibit stage-specific expression patterns (Fig. 9F). The sexual stage had a great number of up-regulated or specifically expressed lncRNAs, making lncRNAs being more prevalent in sexual stage.

**Figure 9.**
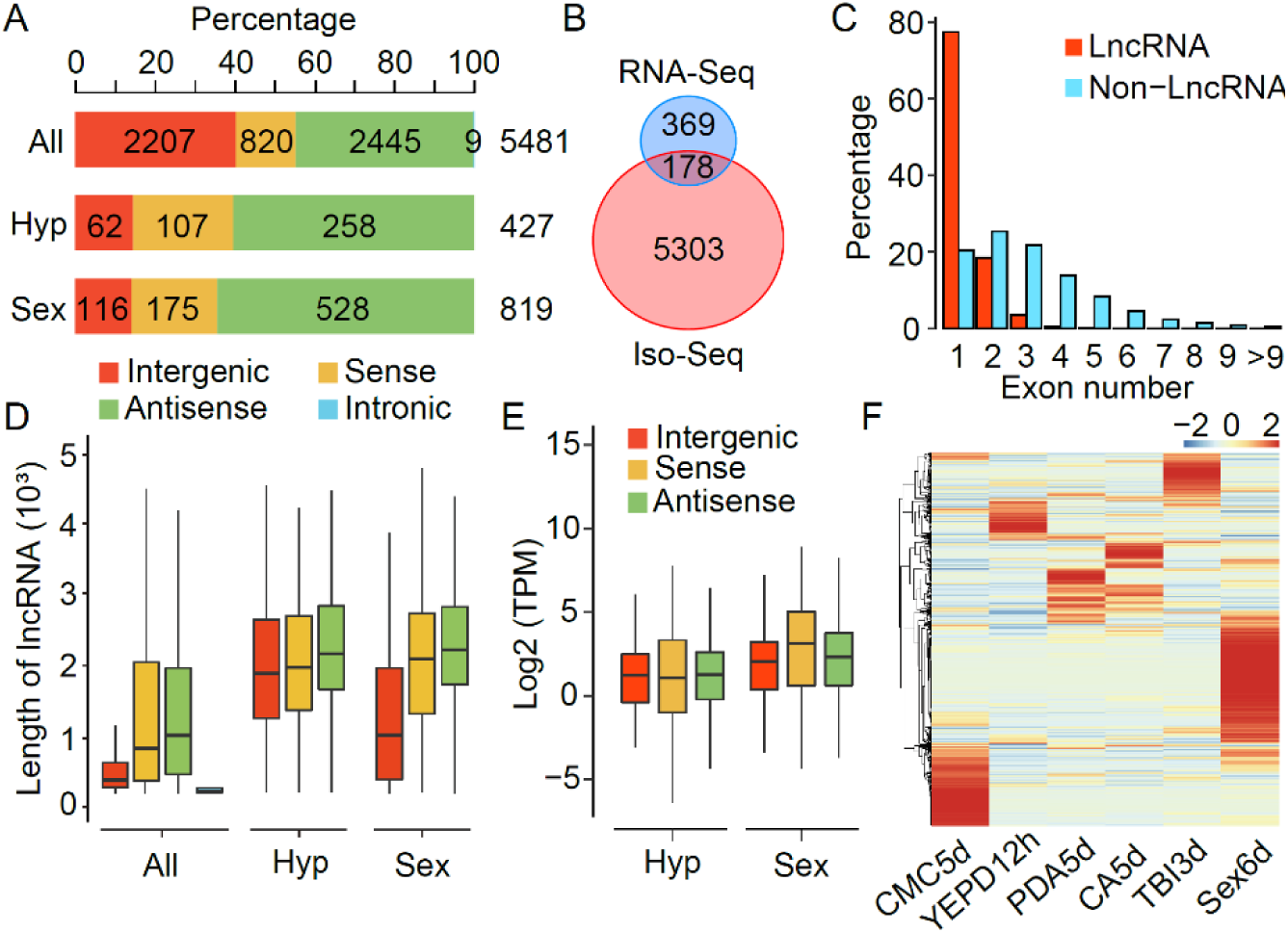
Characterization of lncRNAs in *F. graminearum*. (*A*) Number and percentage of lncRNAs in each category identified in all samples (All) as well as in vegetative hyphae (Hyp) and sexual perithecia (Sex). (*B*) Numbers of distinct and common lncRNAs identified by Iso-Seq in this study and by short-read RNA-Seq in previous study (Kim et al. 2018). (*C*) Distribution of the number of exons in lncRNAs and non-lncRNAs. (*D-E*) Box plots comparing the length of lncRNAs (*D*) and their expression levels (log_2_ TPM) (*E*) in each marked category. (*F*) Heatmap of the expression of lncRNAs across different samples. High (orange to red) and low (yellow to blue) expression levels are depicted as Z-scores for each lncRNA. See Supplemental Table S2 for details of sample information.

### Discovery of polycistronic transcripts in *F. graminearum*

Although polycistronic gene expression, the co-transcription of multiple independently transcribed loci that retain coding potential from one single promoter to generate a polycistronic transcript, was generally considered rare in eukaryotes, widespread polycistronic transcripts were discovered recently in agaricomycete fungi as well as in plant and green algae by using Iso-Seq (Gordon et al. 2015; Wang et al. 2019; Gallaher et al. 2021). Using a similar method (see Methods), we identified 914 polycistronic transcripts that spanned two or more consecutive protein-coding genes or ORFs on the same strand in *F. graminearum* (an example is shown in Fig. 10A). The vast majority (98%) of them were bicistronic transcripts that cover two annotated ORFs (Fig. 10B). Collectively, these polycistronic transcripts were transcribed from 321 loci, corresponding to 4.9% (698) of annotated ORFs. Each polycistronic loci had at least two supporting FLNC reads, suggesting that these transcripts are less likely to represent chimeric reads.

**Figure 10.**
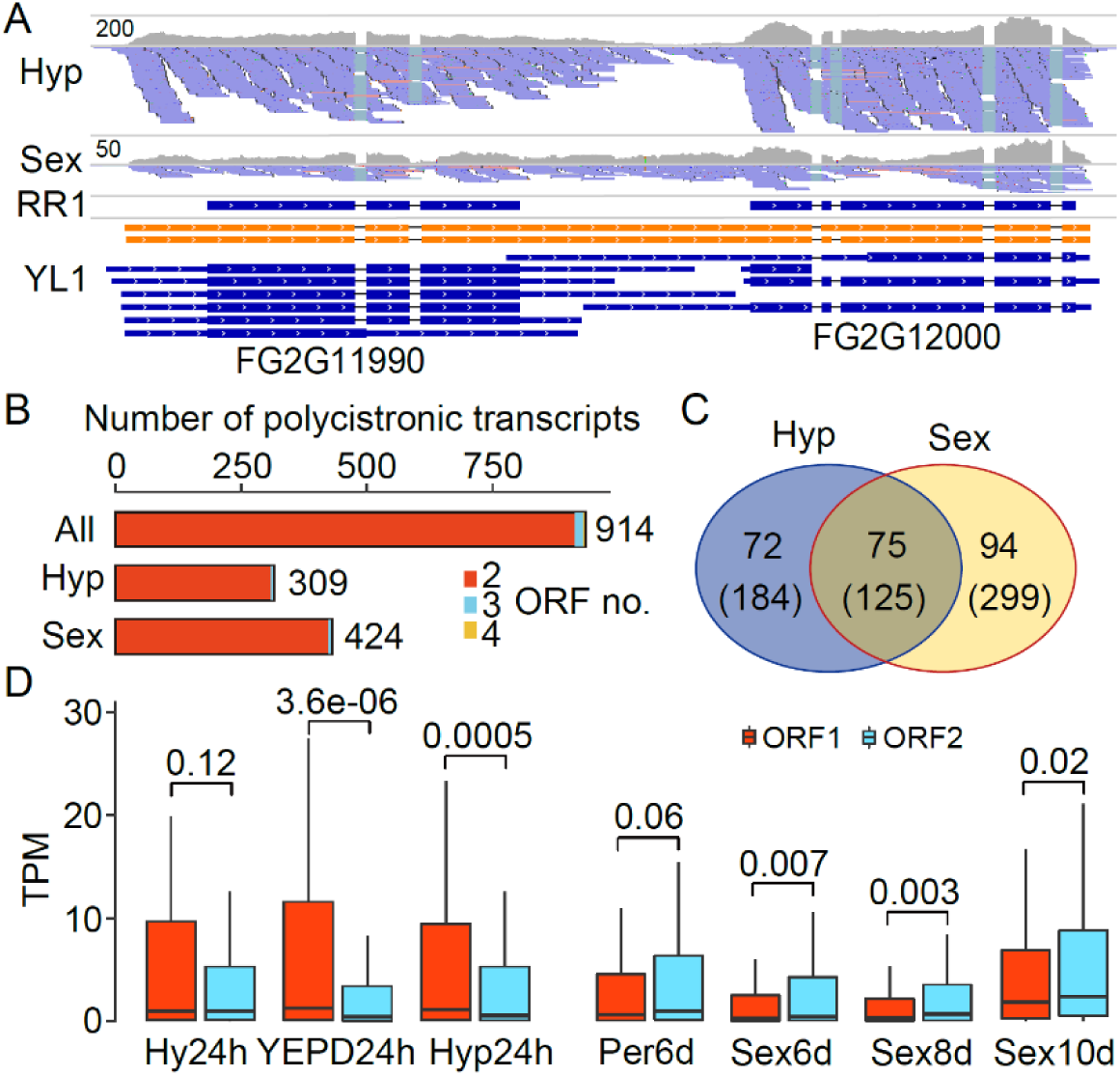
Characterization of polycistronic transcripts in *F. graminearum*. (*A*) An example of polycistronic transcripts spanning two genes supported by Iso-Seq transcripts and RNA-Seq reads. Polycistronic transcripts are shown in orange and non-polycistronic transcripts of each gene (transcribed independently) in blue. (*B*) Number of annotation genes (ORFs) covered by polycistronic transcripts in all samples (All) as well as in vegetative hyphae (Hyp) and sexual perithecia (Sex). (*C*) Numbers of shared or unique polycistronic loci and transcripts (in brackets) between vegetative (Hyp) and sexual (Sex) stages. (*D*) The independent expression levels (TPM) of ORFs within polycistronic transcripts in the marked samples. ORF1 and ORF2 are named after their order in the transcript (5’-to 3’). *P* values are from one-tailed Wilcoxon rank sum test. See Supplemental Table S2 for details of sample information.

A total of 309 and 424 polycistronic transcripts were identified in vegetative growth and sexual stages, respectively (Fig. 10B). Most of them were stage-specific (Fig. 10C), implying that their expression may be under the control of stage specification signals. Like agaricomycete fungi, genes within polycistronic transcripts were also independently transcribed in *F. graminearum* (see the example in Fig. 10A). Interestingly, in comparison with their upstream genes within the same polycistronic transcripts, the expression of downstream genes was generally lower in vegetative growth stage but higher in sexual stage in independent biological replicates (Fig. 10D), suggesting that the expression of downstream genes may be repressed by the upstream readthrough transcription during vegetative growth but induced during sexual reproduction. Therefore, the polycistronic transcripts may play distinct regulatory roles during vegetative growth and sexual reproduction.

## DISCUSSION

The reference genome of *F. graminearum* PH-1 comprising four chromosomes and a mitochondrial genome was originally sequenced by Sanger sequencing (Cuomo et al. 2007) and completed with Illumina resequencing data (King et al. 2015; King et al. 2017). In this study, we further refined the reference genome PH-1 with PacBio SMRT long-read sequencing. To detect spontaneous mutations occurred during culture transfer and maintenance, the Illumina resequencing data of three different lab stocks of PH-1 were generated and used for genome improvement. Totally, 249 error position or regions in the previous genome assembly were corrected. The complete and accurate genomic reference sequence of PH-1 generated in this study is valuable for read mapping and functional studies in *F. graminearum*, and for improving the accuracy and efficacy of genome assembly of closely related species.

Besides accurate genome assembly, high-quality gene and transcript annotation is also critical for comparative and functional genomics. The gene model of PH-1 has been annotated and improved by BROAD institute (Cuomo et al. 2007), Munich Information Services for Protein Sequences (MIPS) (Wong et al. 2011), and Rothamsted Research (RR) (King et al. 2015; King et al. 2017). Although previous gene model annotation was generated by integrating multiple gene prediction algorithms, EST and RNA-Seq data were merely used for gene model support. No genuine transcript information direved from transcript reconstruction was incoporated to the annotation. In this study, we reconstructed the full-length transcriptome of PH-1 with data from PacBio Iso-Seq and deep-depth strand-specific RNA-Seq. Unlike transcriptome reconstruction with RNA from vegetative hyphae only in *A. apis* (Chen et al. 2020), we maximized transcript diversity by sampling six cell types and using multiple size-selected and non-size-selected libraries. Ultimately, a comprehensive reference annotation (YL1) comprising 51,617 transcript isoforms from 17,189 genes was generated for *F. graminearum* PH-1. Compared to the previous annotation (RR1) consisting of only 14,145 transcripts from 14,145 genes, the updated reference annotation YL1 described in this study is much more comprehensive and complete. Majority of the transcript isoforms in YL1 annotation have not been annotated previously. More importantly, a total of 2,998 potential novel protein-coding genes and 5,481 lncRNAs were identified and included in the YL1 reference annotation. To our knowledge, this is the most comprehensive full-length transcriptome documented in filamentous fungi to date, which is a rich resource of transcript isoforms for comparative and functional genomic studies in filamentous ascomycetes. Our methodology for Iso-Seq data analysis will also be a useful reference for analyzing the full-length transcriptome in other fungal species.

Recent studies with short-read RNA-seq data have indicated that the AS frequency in fungi may be substantially underestimated, and up to 63% of the multi-exonic genes in *T. longibrachiatum* (Xie et al. 2015) and 50% in *V. dahliae* (Jin et al. 2017) had AS events. In *F. graminearum*, only 231 genes with AS events were previously reported (Zhao et al. 2013). In this study, we detected a total of 54,613 AS events in 42.7% (4,997) of the multi-exonic genes in *F. graminearum*. One advantage of Iso-Seq is its suitability for identifying not only the AS events but also the splice isoforms. We detected a total of 17,229 splice isoforms in *F. graminearum*. Moreover, we found that 1,960 genes have splice isoforms derived from multiple combinatory AS events. These results suggest that AS significantly increases fungal transcriptome complexity, expanding our view of the regulatory potential of RNA splicing in fungi. Furthermore, detection and quantification analysis of differential AS between sample groups from short-read RNA-Seq data commonly rely on pre-annotation of known spliced transcripts, we discovered massive differential AS events from RNA-Seq data of different cell types in *F. graminearum*, demonstrating the tremendous advantages of our YL1 transcript annotation.

The regulatory function of AS in plants and animals has been well established (Chaudhary et al. 2019). However, the contribution of AS to fungal biology is still elusive. In *Ustilago maydis*, mutants lacking the peroxisomal isoforms of Gapdh or Pgk1 are reduced in virulence (Freitag et al. 2012). Recently, the AS event of *MoPTEN* was reported to be important for growth and pathogenesis in *M. oryzae* (Wang et al. 2021b). In *Sclerotinia sclerotiorum*, a number of AS isoforms are differentially expressed on diverse host plants, which may contribute to its broad host spectrum (Ibrahim et al. 2021). In *F. graminearum*, we found a variety of biological processes were enriched in the genes with AS events, implying that AS may play important regulatory roles. Furthermore, a large number of differential AS events were identified among different cell types, indicating that these AS events may be developmentally regulated in *F. graminearum*. More importantly, we found a global increase in intron inclusion in the aging or dormant tissues relative to the tissues with active hyphal growth, suggesting that IR regulation is associated with aging or dormancy in *F. graminearum*. In animals, recent studies have showed that dysfunctional AS events contribute to the aging/senescence phenotype across multiple species (Deschenes and Chabot 2017; Bhadra et al. 2020; Angarola and Anczukow 2021). A global increase in intron inclusion has been observed in aging tissues of *Caenorhabditis elegans, Drosophila*, mouse, and human (Bhadra et al. 2020; Angarola and Anczukow 2021). In *F. graminearum*, we found that the expression level of spliceosomal genes were altered in conidia and 10-dpf perithecia. Like in animals (Adusumalli et al. 2019), the differentially retained introns in the aging or dormant tissues of *F. graminearum* have a higher GC content although they tend to be longer than the spliced introns. It is tempting to speculate that a global increase in intron inclusion may be a transcriptional signature of aging that is evolutionarily conserved between animals and fungi.

One major role of AS is to generate distinct proteins from the same gene. The contribution of AS toward proteome diversity is well documented in animals and plants (Chaudhary et al. 2019). In *S. cerevisiae* that has only ∼300 intron-containing genes, however, AS is mainly used to control transcript levels rather than generate proteome diversity (Kawashima et al. 2014). In *F. graminearum*, 78.5% of the AS transcript isoforms identified in this study encode proteins with altered sequences relative to canonical transcript isoforms, indicating that AS may significantly increase its proteomic complexity. In fission yeast *Schizosaccharomyces pombe*, 59.7% of translatable isoforms with AS events encode proteins different from the annotated isoforms (Kuang et al. 2017). Similar to *S. pombe*, the predicted alterations in protein sequences due to AS occur more frequently at the beginning of ORFs in *F. graminearum*. However, whether these predicted novel proteins from AS transcript isoforms are actually produced remains to be determined, as multiple proteomic studies suggested that AS does not contribute to proteome complexity as expected in animals and plants (Chaudhary et al. 2019).

Coupling AS to NMD is considered a regulatory mode of gene expression (Garcia-Moreno and Romao 2020). In plants, IR is known to generate mostly AS transcript isoforms harboring PTC (Chaudhary et al. 2019). Because IR is also the predominant AS type in fungi as in plants, we expected most intron-retaining AS transcript isoforms to be potential NMD targets in *F. graminearum*. Unexpectedly, we found only 6.2% of the AS transcript isoforms were predicted as PTC-containing NMD targets in *F. graminearum*. In animals and plants, approximately one-third of all AS isoforms are predicted as PTC-containing NMD targets (Wang et al. 2018; Garcia-Moreno and Romao 2020). More importantly, these PTC-containing NMD targets were not accumulated in the Δ*Fgupf1* mutant. The vast majority of the transcripts up-regulated in the Δ*Fgupf1* mutant were canonical transcript isoforms or AS transcript isoforms without ORF changes. These results suggest that AS may be not coupled to NMD in *F. graminearum*. In the human pathogen *Cryptococcus neoformans*, most intron-retaining transcripts also were not sensitive to NMD (Gonzalez-Hilarion et al. 2016). Even in animals and plants, many PTC-containing transcripts are not subjected to NMD (Chaudhary et al. 2019). In *F. graminearum*, the AS transcript isoforms with shortened ORFs were generally expressed at relatively low levels. Therefore, NMD-independent pathways involving in the clearance of these transcripts may be common in animals, plants and fungi.

Distinct from yeasts, in which depletion of the Upf1 proteins does not affect its growth, the Δ*Fgupf1* mutant had severe growth defects in *F. graminearum*. We detected 672 down-regulated transcripts in Δ*Fgupf1* mutant, which were mostly enriched for GO categories related to ribosome biogenesis. Interestingly, the downregulation of genes involved in translation and ribosome biogenesis was also found in the *upf1* mutant of Arabidopsis recently (Raxwal et al. 2020). It is most likely that the FgUpf1 play an important role in translational gene regulation in *F. graminearum* as suggested in Arabidopsis.

Pre-mRNA 3’-end processing have been studied extensively in yeasts and animals, but rarely in filamentous fungi (Vavasseur and Shi 2014; Tian and Manley 2017; Gruber and Zavolan 2019). APA plays important regulatory roles in virulence and development in *M. oryzae* (Franceschetti et al. 2011). Using a short-read sequencing-based 3’ T-fill method, 14,593 PASs were identified and 52% (4,283) of *M. oryzae* genes were found to be alternatively polyadenylated (Rodriguez-Romero et al. 2019). PASs can be accurately identified from the Iso-Seq FLNC reads at single-nucleotide resolution. We identified 364,513 unique PASs and showed that 64.8% (11,133) of genes had more than one PASs in *F. graminearum*, excluding microheterogeneity. These results suggest that APA is pervasive in filamentous fungi as in mammals and plants (Wu et al. 2011; Tian and Manley 2017). Moreover, four conserved sequence motifs were identified surrounding the PASs and they were more enriched at distal PASs compared to proximal PASs. Therefore, distal PASs may be stronger than proximal ones in *F. graminearum*, which is similar to *S. pombe* and mammals but in contrast to *S. cerevisiae* (Liu et al. 2017; Gruber and Zavolan 2019). However, the proximal PAS isoforms generally had higher expression levels than the distal PAS isoforms in *F. graminearum*, which is similar to what has been reported in *S. cerevisiae* but opposite to observations in *S. pombe* and mammals (Hoque et al. 2013; Liu et al. 2017). This raises the question of why the weaker proximal PASs are preferentially used in *F. graminearum*.

Core mRNA 3’-end processing factors are known to affect the processing of particular PASs (Gruber and Zavolan 2019). By performing gene knockout and strand-specific RNA-Seq, we showed that the core 3’-end processing factors FgRNA15, FgHRP1, and FgFIP1 all play important roles in recognizing the sequence motifs and promoting proximal PAS usage in *F. graminearum*. Deletion of them resulted in global increase in distal PAS usage. These results are consistent with the findings in humans, in which the core 3’-end processing factors CSTF2/CSTF64, FIP1L1, and CFIm that are homologous to yeast RNA15, FIP1, and HRP1, respectively, promote cleavage and polyadenylation at proximal PASs (Gruber and Zavolan 2019; Pereira-Castro and Moreira 2021). However, unlike CFIm in humans, FgRNA15 has a particularly strong impact on PAS selection in *F. graminearum*. Furthermore, we found increased intron retention in these three mutants, especially in the Δ*Fgrna15* mutant, suggesting that the three 3’-end processing factors may also promote intron splicing in *F. graminearum*. A role for 3’-end processing factors in promoting AS was reported recently in humans (Misra et al. 2015), where the CPSF (homologous to the fungal CPF: CFII subcomplex) and SYMPK, but not other 3’-end processing factors, have a global role in promoting both inclusion and exclusion of internal exons. Our study indicates that the subunits of different 3’-end processing complex/subcomplex could be involved in promoting AS in fungi, including CFIA (FgRNA15), CFIB (FgHRP1), and CPF: PFI (FgFIP1).

Since 3’-UTR affects the stability, translation rate, and subcellular localization of mRNAs (Pereira-Castro and Moreira 2021), changes in the ratio of 3’-UTR isoforms in different cell types may affect cellular functions of corresponding proteins. In this study, we found a global increase in distal PAS usage in dormant conidia and old tissues compared to younger tissues in *F. graminearum*. Moreover, genes with increased distal PAS usage in the dormant and old tissues were enriched for the ones functionally related to many senescence/age-related pathways as reported in mammals (Deschenes and Chabot 2017; Chen et al. 2018; Angarola and Anczukow 2021). These results suggest that APA-mediated 3’-UTR lengthening may play a role in regulating aging and dormancy in *F. graminearum*. In mammals, short 3’-UTR isoforms are usually associated with cell proliferation and activation while long 3’-UTR isoforms are mostly found in polarized and differentiated cells (Pereira-Castro and Moreira 2021). Global lengthening of 3’-UTRs in senescent cells was only described recently (Chen et al. 2018). In yeasts, the expression of long 3’-UTR isoforms is favored in nutrient-restricted conditions, in which cells reaching the quiescent state (Liu et al. 2017). In mammals, 3’- end processing factors play important roles in APA regulation. It has been proposed that the generation of short 3’-UTR in proliferating cells is due to global upregulation of the 3’-end processing machinery. In *F. graminearum*, the core 3’-end processing factors were generally downregulated in the older or dormant tissues, suggesting that the increase of distal PAS usage in generating long 3’-UTR isoforms in the older or dormant tissues is likely due to global downregulation of the 3’-end processing machinery. Therefore, APA contributes to the regulation of gene expression during cell senescence/aging and dormancy may be evolutionarily conserved in mammals and fungi.

In animal cells, short 3’-UTR isoforms were generally more stable than long 3’-UTR isoforms, and both long 5’- and 3’-UTR isoforms tended to translate less efficiently (Zhou et al. 2019; Pereira-Castro and Moreira 2021). Retained introns located in the 5’- or 3’-UTR may increase the lengths of affected UTRs. Therefore, our findings of the global increase in both intron inclusion and distal PAS usage in aging and dormant tissues demonstrate that AS and APA as an integral part of cell reprogramming processes to regulate the overall protein production in *F. graminearum*.

In *F. graminearum*, 547 lncRNAs identified by short-read RNA-Seq analysis are under stage-specific regulation during fruiting body formation (Kim et al. 2018). In this study, we identified 5,481 high-confidence lncRNAs from Iso-Seq data, which exhibited stage-specific expression patterns. In addition, we discovered 914 polycistronic transcripts that were found to be expressed in a stage-specific manner. Particularly, the expression level of downstream genes covered by the polycistronic transcripts was lower in vegetative growth stage but higher in sexual stage, implying that the polycistronic transcripts may play distinct regulatory roles in vegetative growth and sexual stages. The lncRNAs and polycistronic transcripts identified here provide fundamental resources for further investigations on their roles in fungal biology.

Overall, this study represents the first large-scale analysis of a full-length transcriptome in a filamentous plant pathogenic fungus, providing new insights into the complexity and regulation of fungal transcriptomes. The updated genome sequences and comprehensive reference set of transcript isoforms generated in this study will be beneficial to the *Fusarium* community and fungal community in general for comparative and functional genomic studies.

## METHODS

### Fungal materials and growth conditions

The wild-type *F. graminearum* strain PH-1 (Cuomo et al. 2007) maintained in our lab (PH-1_YL) and its mutants generated in this study were routinely cultured on potato dextrose agar (PDA) plates at 25°. The PH-1 lab stocks maintained at two other labs were kindly provided by Dr. Frances Trail at Michigan State University (PH-1_MSU) and Dr. Yun Chen at Zhejiang University (PH-1_ZJU), respectively. For sample collection, fungal tissues at different developmental stages or conditions were harvested as described (Liu et al. 2015; Liu et al. 2016). Briefly, perithecia were harvested from carrot agar (CA) plates at 3-, 6-, 7- or 8-dpf. Conidiation samples were collected from 5-day-old liquid carboxymethyl cellulose (CMC) cultures. Conidia were collected by filtering 5-day-old CMC cultures through sterile glass wool. Germlings and vegetative hyphae germinated from conidia were collected from 12, 24, 36, and 48 h liquid YEPD (1% yeast extract, 2% peptone, 2% glucose) cultures, respectively. Aerial mycelia were collected from 5-day-old PDA and CA plates, respectively. DON-producing hyphae were collected from 3-day-old liquid trichothecene biosynthesis induction (TBI) cultures supplied with 5 mM arginine or NH_4_NO_3_. The inoculated spikelets of flowering wheat heads of cultivar Xiaoyan22 were collected 3 days after inoculation (dpi) with PH-1. Samples were immediately frozen in liquid nitrogen.

### Generation of gene deletion mutants

The split-marker approach (Catlett et al. 2003) was used to generate the gene replacement constructs for the *FgCLP1, FgFIP1, FgHRP1, FgRNA15, FgUPF1* and *FgYTH1* genes. The flanking sequences of individual genes were amplified and connected to the hygromycin phosphotransferase (hph) cassette by overlapping PCR with the primers listed in Supplemental Table S6. Protoplasts of the PH-1-YL1 strain were prepared and transformed with each gene replacement construct as described previously (Liu et al. 2015). For transformant selection, hygromycin B (Calbiochem, La Jolla, CA) was added to the final concentration of 250 mg/ml. Gene deletion mutants were confirmed by PCR assays. At least two independent deletion mutants were obtained for each gene.

### PacBio and Illumina DNA library preparation and sequencing

Genomic DNA was isolated from 24 h vegetative hyphae as described (Murray and Thompson 1980). Quality of DNA was evaluated by agarose gel electrophoresis. PacBio library was constructed using the SMRTbell^™^ Template Prep Kit version 1.0 according to manufacturer’s instructions, and sequenced on the PacBio Sequel instrument. Illumina library was prepared with the NEBNext^®^ Ultra™ DNA Library Prep Kit for Illumina^®^ following the manufacturer’s instruction and sequenced on the Illumina HiSeq^®^ 2500 System, with a 2×150 bp paired-end read mode.

### PacBio Iso-Seq library preparation and sequencing

Total RNA was extracted and purified with the RNAprep Pure Plant Kit (Tiangen Biotech, Beijing, China). Poly(A)^+^ mRNA was enriched with oligo (dT) magnetic beads and analyzed with BioAnalyzer-2100 (Agilent). Purified poly(A)^+^ mRNAs from samples of CA5d, CMC5d, PDA5d, Sex6d, TBI3d, YEPD12h were pooled together in equimolar ratios for subsequent library construction. The Iso-Seq library was prepared according to the Iso-Seq™ Template Preparation for Sequel^™^ Systems protocol. In brief, RNA is synthesized to cDNA using the Clontech SMARTer PCR cDNA Synthesis Kit, and subsequently amplified to generate double-stranded cDNA. The cDNA is then constructed to a SMRTbell library for sequencing with the PacBio Template Prep Kit. For the size selected Iso-Seq libraries of 1-2, 2-3, and 3-6 kb, the amplified cDNA product was size selected with the BluePippin before library construction. The SMRTbell libraries was sequenced on the PacBio RS II or Sequel System.

### Illumina strand-specific RNA-Seq library preparation and sequencing

Total RNA was extracted and purified with the RNAprep Pure Plant Kit (Tiangen Biotech, Beijing, China) from each sample list in Supplemental Table S2. The total RNAs of the Coni, DON3d, Ger12h, Hyp24h, Inf3d, Sex3d, Sex8d and Sex10d samples were subject to ribosomal RNA (rRNA) depletion using the Ribo-Zero rRNA Removal Kit (Illumina, USA) according to the manufacturer’s protocol. The poly(A)^+^ mRNAs from other samples were enriched using oligo (dT) magnetic beads. Quality of RNA was evaluated by agarose gel electrophoresis and BioAnalyzer-2100 (Agilent). Strand-specific RNA-Seq libraries were prepared with the NEBNext^®^ Ultra^™^ Directional RNA Library Prep Kit following the manufacturer’s instruction, and sequenced using the Illumina HiSeq^®^ 2500 system with the 2×150 bp paired-end read mode.

### Reference genome correction

The public genome assembly (RR1, release-35) of PH-1 was download from Ensembl Fungi. PacBio subreads from SMRT sequencing were obtained by using SMART link v5.1.0 (Gordon et al. 2015) with the default setting. Only subreads longer than 1 kb were kept as high-quality reads and used for further analyses. The *de novo* genome assembly was generated with *Canu* software version 1.7 (Koren et al. 2017) using default parameters and polished with *Arrow* from SMRT link software. The PacBio genome assembly was aligned to the RR1 assembly using *mummer4* (Marcais et al. 2018), and different sites or regions between them were identified by our in-house Python script based on the alignments. The PacBio reads and the Illumina reads of

PH-1 lab stocks from three different labs (PH-1_YL, PH-1_MSU, and PH-1_ZJU) were mapped onto the two assemblies by *GMAP* (Wu and Watanabe 2005) and *Bowtie2* (Langmead and Salzberg 2012), respectively. Each site or region that differ between the two assemblies were checked manually by using IGV (Robinson et al. 2011). Mis-assembly, base and indel errors in the RR1 assembly were identified and corrected based on the alignments. Only the errors evidenced by at least two sequenced PH-1 lab stocks were corrected.

### Iso-Seq analysis pipeline

Our Iso-Seq analysis pipeline consists of six steps (Supplemental Fig. S1). ***Step1: Generating polished consensus transcript isoforms***. The *pbtranscript* tool included in SMRT Link (Gordon et al. 2015) was used to extract the Reads of Insert (ROI) from raw Iso-Seq datasets generated by SMRT sequencing, classify them into full-length, non-artificial-concatemer (FLNC) and non-full-length, non-artificial-concatemer (nFLNC) reads based on the location of primers and poly(A) tails, and then cluster and polish the FLNC reads into polished consensus transcript isoform sequences. ***Step2: Correcting indels, mismatches and splice junction using genome sequence and strand-specific RNA-Seq***. First, the RNA-Seq data from the same RNA as Iso-Seq was used to generate the splicing junction file (SJ.out.tab) using *STAR* (Dobin et al. 2013) with parameter --twopassMode Basic. Then, the consensus transcript isoform sequences were repaired using *TranscriptClean* (Wyman and Mortazavi 2019) according to the genome sequence and splicing junction file. ***Step3: Detecting and filtering fusion and polycistronic transcripts***. Because the presence of fusion or polycistronic transcripts can disturb subsequent gene (transcription unit) definition by the Cupcake script *collapse_isoforms_by_sam*.*py*, they were detected and filtered out from the consensus transcript isoform sequences. Putative fusion transcripts were identified using the Cupcake script *fusion_finder*.*py*. Polycistronic transcripts were detected according to the standard that a polycistronic transcript must have two or more non-overlapping ORFs (≧100 aa) and each ORF overlaps over 50% CDS region of corresponding annotated genes. ***Step4: Filtering artificial-concatemer transcripts and correcting wrong-stranded transcripts***. The polished consensus transcript isoform sequences were aligned to the reference genome by *BLAT* (Kent 2002), and the artificial-concatemer transcripts were identified and filtered out according to their mapping characteristics by custom Python script. To detect wrong-stranded transcripts, the polished consensus transcripts were first aligned to the reference genome and then converted to GFF format using *GMAP* (Wu and Watanabe 2005). The transcripts that aligned to the annotated genes in opposite strand but shared the same splicing junction were identified by using *GffCompare* (Pertea et al. 2016) and corrected for their strand information. ***Step5: generating unique transcript isoform annotation***. The filtered and corrected consensus transcript isoform sequences were mapped to the reference genome using *GMAP*. Based on the mappings, redundant transcript isoforms were collapsed into unique transcript isoforms by the Cupcake script *collapse_isoforms_by_sam*.*py* (min-coverage=0.9 and --min-identity=0.9). Since the Iso-Seq cDNA library protocol does not guarantee transcript isoform sequences that preserve the 5’ start, to minimize inclusion of possible 5’ truncated transcripts, transcript isoforms differing only in the 5’ start of their first exon were collapsed to keep only the longest ones. To avoid filtering out some real transcript isoforms with alternative transcription start sites, we therefore generated a 5’ start un-collapsed transcript isoform annotation file available in the FgBase although only the collapsed transcript isoform annotation was used in this study. Additionally, genes in the transcript isoform annotation that only have one single-exon transcript with short length (≤200 nt) and low expression (≤1 TPM) were removed. ***Step6: Correcting unique transcript isoform annotation***. Because of high gene density in *F. graminearum*, transcripts from adjacent genes in same strand commonly overlapped. Overlapping transcripts were grouped as one gene by Cupcake script. To correct these issues, the unique transcript isoform annotation was compared with the RR1 gene annotation by *GffCompare*. The genes from transcript isoform annotation that overlap with two or more RR1 genes were checked and corrected manually with the aid of IGV.

### RNA-Seq data analysis

RNA-Seq data were aligned to the reference genome using *HISAT2* (Pertea et al. 2016). Transcript assembly and quantification of gene and transcript expression were performed using *StringTie* (Pertea et al. 2016). TPM (Transcripts Per Kilobase Million) is used as the normalized unit of gene or transcript expression. Differential expression was analyzed with *cuffdiff* (Trapnell et al. 2013). Only genes or transcripts with a minimum expression level of 1 TPM in at least one RNA-Seq library were included in the differential expression analysis. Genes or transcripts with a q_value≤0.05 and |log_2_ fold change|≥1 were considered to be differentially expressed.

### AS events identification and analysis

AS landscape in the YL1 reference annotation datasets were extracted using *AStalavista* web server (astalavista.sammeth.net/). According to the AS code assigned, the AS events were categorized as four basic types with “(n)^, (n+1)-” for IR, “(n)^,(n+1)^” for A5, “(n)-, (n+1)-” for A3, and “(n)-, (n+1)^” for ES. Differential AS events among samples were identified by *CASH v2*.*2*.*1* (Wu et al. 2018) based on the YL1 annotation. AS events with a false discovery rate (FDR) < 0.05 were regarded as differential AS events.

### NMD target prediction

The position and distance of the stop codon relative to the terminal exon-exon junction was determined for each AS transcript isoforms. AS transcripts with a stop codon >50-bp upstream of the terminal exon-exon junction were considered as putative NMD targets (Garcia-Moreno and Romao 2020). The PTC-containing NMD targets were identified by comparation of the stop codon position of the predicted NMD target with that of the corresponding canonical transcript.

### APA identification and analysis

The FLNC reads were aligned to the YL1 genome and used to identify the unique PASs. To address the internal priming issue, the PASs with AAAAAA in the upstream 10 nt and downstream 20 nt or with more than 7 As in 10 nt sliding windows were discarded. For each confident PASs, the number of supporting FLNC reads were counted. Separating APA from heterogeneous cleavage was performed with a similar method as described (Liu et al. 2017). In brief, the expectation maximization (EM) algorithm in the *mixtools* package of *R* programming (Benaglia et al. 2009) was used to identify two distribution modes based on the distance between adjacent PASs. This method resulted in a cross point of ∼9 nt between the two distribution modes. The adjacent PASs with a distance >9 nt were separated into two PAS cluster, whereas those ≤ 9 nt were clustered together as one PAS cluster.

The PAS with the highest number of supporting FLNC reads in a PAS cluster was chosen as the representative site for the cluster. *SignalSleuth2* (Zhao et al. 2014) and *MEME-chIP* (Machanick and Bailey 2011) were used to identify enriched 4-mer to 6-mer motifs in flanking sequences (−100 nt to 100 nt) around the PASs. Detecting APA and identifying differential APA usage from RNA-seq alignments were performed with the *roar* software (Grassi et al. 2016). Since the proximal PAS isoform generally had a higher expression level than the distal PAS isoform in *F. graminearum*, we used the M/m value in this study by taking the inverse of the m/M value reported by *roar* (Grassi et al. 2016). PAS with a p-value ≤ 0.05 was considered to be differential APA usage.

### LncRNA identification

Novel transcripts from Iso-Seq transcript isoform set that did not share any splice junctions with RR1 genes were used to predict lncRNAs. We used *PLEK* (Li et al. 2014) to distinguish lncRNAs from protein-coding RNAs. The transcripts with a low coding potential were further scanned against the Pfam and Rfam databases to filter out transcripts encoding protein domains and/or harboring any known structural RNA motifs (E value <10^10^).

### RT-PCR

Primers used for PCR validation of AS events were designed to span the splicing events using *Primer Premier 5*. RNA samples were isolated with the TRIzol reagent (Invitrogen, Carlsbad, CA, USA) from 24 h vegetative hyphae and 6-dpf perithecia. The ReverAid First cDNA synthesis kit (Thermo Fisher Scientific) was used for cDNA synthesis. PCR products were purified and sequenced by Sanger method.

### Statistical analysis

All statistical tests were performed using *R* (www.r-project.org). Hierarchical clustering analysis was performed using *R* with average linkage cluster method and the Pearson correlation coefficient as the distance value. Heatmaps were generated by the package *pheatmap*. GO enrichment analysis was performed using BLAST2GO (www.blast2go.com) with the Fisher’s Exact Test and Benjamini-Hochberg correction (FDR<0.05). Only the most specific GO terms were retained.

### DATA ACCESS

All raw sequencing data generated in this study have been submitted to the NCBI Sequence Read Archive (SRA) under accession numbers listed in Supplemental Table S1 and S2. The YL1 reference genome and transcript annotation are freely available in the customized genome database FgBase (fgbase.wheatscab.com).

## ACKNOWLEDGEMENTS

We thank Mengchun Wu for preparing the fungal samples, Novogene for preparing the DNA and RNA libraries and sequencing; Drs Xue Zhang, Guanghui Wang, and Chenfang Wang for helpful discussions; and Drs Frances Trail and Yun Chen for kindly providing the PH-1 lab stocks. This study was supported by funding for HL from the National Key R&D Program of China (2019YFD1000605), National Natural Science Foundation of China (no. 31872918), National Youth Talent Support Program (Z111021802), and the Programme of Introducing Talents of Innovative Discipline to Universities from the State Administration of Foreign Experts Affairs (no. B18042), and funding for JRX from USWBSI.

## AUTHORS’ CONTRIBUTIONS

HL and PL designed the experiments and analysis pipelines. DC, ZQ, and YC performed the experiments, PL, HW, and QW performed the analyses. HL, JRX, and CJ contributed advice and reagents. The figures were prepared by PL, HW, and HL. The manuscript was written by HL, PL, and JRX. All authors read and approved the final manuscript.

